# Measuring prion propagation in single bacteria elucidates mechanism of loss

**DOI:** 10.1101/2023.01.11.523042

**Authors:** Krista Jager, Maria Teresa Orozco-Hidalgo, Benjamin Lennart Springstein, Euan Joly-Smith, Fotini Papazotos, EmilyKate McDonough, Eleanor Fleming, Giselle McCallum, Andreas Hilfinger, Ann Hochschild, Laurent Potvin-Trottier

## Abstract

Prions are self-propagating protein aggregates formed by specific proteins that can adopt alternative folds. Prions were discovered as the cause of the fatal transmissible spongiform encephalopathies in mammals, but prions can also constitute non-toxic protein-based elements of inheritance in fungi and other species. Prion propagation has recently been shown to occur in bacteria for more than a hundred cell divisions, yet a fraction of cells in these lineages lost the prion through an unknown mechanism. Here, we investigate prion propagation in single bacterial cells as they divide using microfluidics and fluorescence microscopy. We show that the propagation occurs in two distinct modes with distinct stability and inheritance characteristics. We find that the prion is lost through random partitioning of aggregates to one of the two daughter cells at division. Extending our findings to prion domains from two orthologous proteins, we observe similar propagation and loss properties. Our findings also provide support for the suggestion that bacterial prions can form more than one self-propagating state. We implement a stochastic version of the molecular model of prion propagation from yeast and mammals that recapitulates all the observed single-cell properties. This model highlights challenges for prion propagation that are unique to prokaryotes and illustrates the conservation of fundamental characteristics of prion propagation across domains of life.

---

Prion-forming proteins (hereafter prion proteins) are proteins that can adopt multiple conformations, of which at least one is self-propagating. Prions were originally discovered as the cause of devastating neurodegenerative diseases, such as Creutzfeldt-Jakob’s disease (CJD), in mammals (1). Subsequently, non-pathogenic prions were found across diverse species – such as budding yeast (2–6), *Drosophila* (7), *Arabidopsis* (8), and mammals (9–11) – where they are thought to function as protein-based carriers of epigenetic information. In many cases, the prion capability is conferred on the protein by a modular prion domain (PrD), necessary and sufficient for formation of the prion. Conversion from the soluble form to the prion form (a highly structured aggregated form in many well-studied cases) bestows a loss-of-function (12) or gain-of-function (10, 13, 14) to the attached protein, which can result in a fitness advantage under certain environmental conditions (4–6, 15, 16). A particular property of prions is that they can sometimes form multiple structures, called strains, each of which propagates itself with different properties. In mammals, different strains of the prion protein (PrP) are the cause of different diseases (17, 18), while in yeast different strains of the intensively studied prion [*PSI*^+^] (formed by the essential translation release factor Sup35) differ in their stabilities and aggregate size distributions (19–21).

While the detailed molecular mechanisms of prion propagation are under investigation (22, 23), studies in yeast and mammals appear to be consistent with the nucleated polymerization model (24–26). In this model, proteins are converted from the soluble form to the prion form by elongation of existing oligomeric prion aggregates, while aggregates can be fragmented into smaller oligomers (presumably by chaperones like Hsp104, an ATP-dependent disaggregase that is required for prion propagation in yeast (27)). Initial conversion to the prion form is suggested to happen by the rare spontaneous oligomerization to a critical size *n*, below which oligomers would revert to the soluble form.

Recently, thousands of candidate prion domains (cPrDs) have been identified in bacteria using bioinformatic analyses (28). So far, two of these domains were found to form self-propagating prion aggregates in *Escherichia coli*: the PrDs from the Rho termination factor of *Clostridium botulinum* (*Cb* Rho, (28)) and from the single-stranded DNA binding protein of *Campylobacter hominis* (*Ch* SSB, (29)). Of note, many orthologs of these proteins also have predicted cPrDs (28). Although individual lineages could propagate the prions for more than a hundred generations, a fraction of the cells in each lineage was seen to have lost the prion at each replating round (28, 29). The mechanisms by which the prion is lost, and how long individual cells propagate the prion, are unknown. In the previous study of the *Ch* SSB PrD, two types of lineages were observed, one exhibiting a high-stability phenotype and one exhibiting a lower-stability phenotype, suggesting that prion strains could also exist in bacteria (29). In addition, although the molecular mechanisms of prion propagation appear conserved across mammals and yeast, it is unknown if this apparent conservation of mechanism also extends to bacteria.

In this study, we sought to address these questions by measuring prion propagation in single bacteria. Using microfluidics, single-cell time-lapse microscopy, and mathematical modeling, we uncover how the *Ch* SSB PrD prion (hereafter the *Ch* SSB prion) is propagated and lost. We find that the prion is propagated in two distinct modes with aggregates of different size and stability. We discover that the loss of the prion was caused mainly by stochastic inheritance of the aggregates to only one of the two daughter cells at division (i.e. “partitioning errors”). We show that two orthologous SSB cPrDs also form self-propagating prion aggregates, and that the modes of propagation and loss are conserved in these domains. In addition, we describe lineage-specific differences in the stabilities of prion propagation, thus providing additional support for the previous suggestion that bacterial prions, like yeast prions, can exist as phenotypically distinct strains. We also describe a *Ch* SSB PrD mutant that undergoes conversion to the prion form more readily than the wild-type domain. We implement a stochastic version of the nucleated polymerization model, which strikingly recapitulates all the observed single-cell properties. We use this model to further corroborate our finding that prion loss is caused by partitioning errors by making a prediction, which we then validate experimentally. The model also allows us to estimate the prion replication rate, which is found to be similar to that of mammalian prions. This work provides a new assay for studying prion propagation in individual cells, provides insights on prion propagation and loss, and further establishes the conservation of prion propagation mechanisms across domains of life.

## Results

### Experimental system to track prion propagation and loss in single cells

To investigate how long individual cells propagate a prion and the mechanisms of prion loss, we developed an experimental system that enables us to track prion propagation in thousands of individual cells for many cell divisions (Fig. 1a-d). For this, we used the previously constructed His6-mEYFP-*Ch* SSB-PrD (hereafter *Ch* SSB PrD) fusion protein (29) to visualize prion propagation using fluorescence microscopy. Like the Sup35 prion protein in yeast (30–32), *Ch* SSB PrD requires the presence of a pre-existing prion known as [*PIN*^+^] (for [*<u>P</u>SI*^+^] <u>in</u>ducibility) to access the prion conformation, but not for its maintenance (i.e. the propagation phase) (29). Several prion proteins can serve as [*PIN*^+^], including the *Saccharomyces cerevisiae* New1 protein (29, 30, 32, 33). Therefore, to study prion propagation, we transiently expressed a New1-mScarlet-I fusion on a temperature-sensitive plasmid. After inducing synthesis of the New1 fusion protein and subsequently curing the cells of the New1-encoding plasmid (verified by antibiotic sensitivity and absence of mScarlet-I signal, Fig. S1a-c), colonies containing prion-propagating cells were identified using a previously developed reporter system (29). Specifically, cells containing prion aggregates were previously shown to have elevated levels of the chaperone ClpB, such that colonies containing such cells can be distinguished on X-gal-containing plates using a P*clpB-lacZ* reporter (29). As expected, dark blue colonies displayed visible protein aggregation of the *Ch* SSB PrD (as observed by fluorescence microscopy) in a fraction of the cells, and cell extracts prepared from blue colony cultures contained characteristic SDS-stable aggregates (as observed by semi-denaturing detergent agarose gel electrophoresis; SDD-AGE) (Fig. 1b-c). In contrast, the cells in pale blue colonies showed diffuse fluorescence and contained no SDS-stable protein aggregates (Fig. 1b-c). As previously observed (29), replating dark blue colonies gave both dark and pale colonies, while replating pale colonies resulted in only pale colonies. We thus concluded that dark blue colonies contain a mixture of cells with self-propagating prion aggregates displaying aggregated fluorescence and cells with the protein in the soluble form exhibiting diffuse fluorescence.

**Fig. 1.**
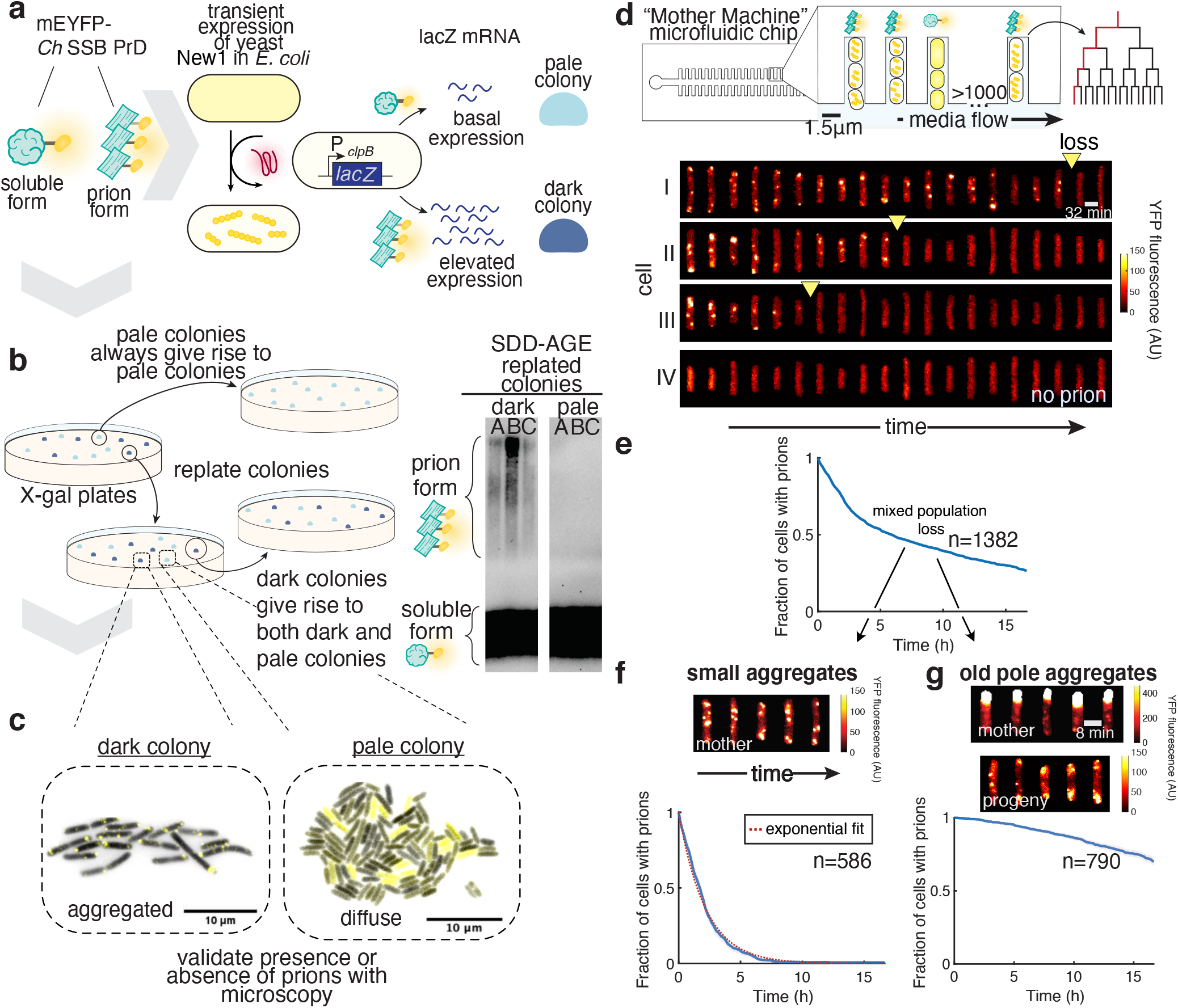
Experimental setup enables quantification of prion dynamics in single cells. **a)** Transient expression of the *S. cerevisiae* New1 protein induces conversion of His6-mEYFP-Ch SSB PrD from its soluble form into the prion form in *E. coli*. Bacteria with prions have elevated levels of ClpB, such that bacterial colonies with prion-containing cells can be distinguished from colonies with cells containing the protein in the soluble form using a P*clpB-lacZ* transcriptional reporter (dark blue vs pale colonies, respectively). **b)** Dark blue colonies contain self-propagating aggregates. (Left) Replating dark blue colonies results in a mix of dark and pale colonies, while replating pale colonies results in only pale colonies. (Right) SDD-AGE shows that different dark blue colonies (A, B and C) contain SDS-stable aggregates, whereas pale colonies contain only soluble *Ch* SSB PrD (prion formation was induced with New1-CFP; a gel where induction was done with New1-mScarlet-I can be found in Fig. S1d). **c)** Fluorescence microscopy images of *E. coli* expressing His6-mEYFP-Ch SSB PrD shows that cells from dark colonies display visible fluorescence aggregation, whereas cells from pale colonies display diffuse YFP fluorescence. **d)** After prion conversion, cells from a dark blue colony are loaded in a microfluidic device where cells are trapped in dead-end trenches and newborn cells are washed away by the flow of growth medium. Fluorescence time-lapse microscopy montage (kymographs) of individual lineages shows that cells propagate the aggregates for heterogeneous duration (I-III) before irreversibly reverting to diffuse fluorescence. YFP fluorescence is shown false-colored according to the colormap indicated on the graph. The prion loss called by our spot-finding algorithm is indicated by a yellow triangle. Cells that have diffuse fluorescence at the beginning of the experiments maintain it (IV). **e)** The fraction of cells with prions over time (prion loss curve) for all aggregate phenotypes shows a biphasic decay, suggesting the presence of two distinct subpopulations. **f)** The prion loss curve for cells with small aggregates fits well to an exponential distribution (red line, R = 0.92). Representative kymograph of cells with small aggregates (top) **g)** Loss curve for cells with old-pole aggregates. Kymographs for the tracked cell (mother) and its progeny (top). The old-pole aggregate is mostly immobile, and the progeny contain small aggregates. The colormap for the old-pole aggregate is different as these aggregates are brighter. The standard error on the mean (SEM) in **e-g** was estimated by bootstrapping, and an envelope is shown as 2xSEM.

For time-lapse microscopy, cells from a single colony containing prion-propagating cells were loaded into a microfluidic device (34) where cells are trapped in short trenches and the newborn cells are washed away by the constant flow of growth media (Fig. 1d). Automated time-lapse microscopy and analysis enables us to track individual lineages for more than two dozen cell divisions while precisely measuring cell fluorescence, growth rate, size, and other characteristics. Using this approach, we observed that cells propagated the prion (aggregated fluorescence, Fig. 1d) over multiple cell divisions before irreversibly losing the prion (diffuse fluorescence, Fig. 1d). Even though the protein concentration was constant throughout the experiment (after reaching equilibrium of growth conditions, SI Materials and Methods 2.5.3.2, Fig. S2a-c), individual lineages displayed remarkable variation in the duration of prion propagation; some cells lost the prion after a few divisions while others kept it for the whole duration of the experiment (~30 divisions).

### Prion propagation occurs through two distinct modes

We next sought to quantify how long individual cells could maintain the prion. For the analyses, we define the time of prion loss as the last time aggregates were detected using a spot-finding algorithm (SI Materials and Methods 2.5.3.3.1, Fig. S3a-c). Counting the detectable aggregates showed that aggregates were both lost and generated until the irreversible loss event, supporting the idea that the prion is propagated during the experiment rather than being simply diluted (Fig. S3d). To measure the distribution of propagation duration, we calculated the fraction of tracked cells containing prion aggregates as a function of time (SI Materials and Methods 2.5.3.3.2). We observed a loss curve with two phases: an initial phase of rapid loss followed by a phase with a slower rate of loss (Fig. 1e). This result suggested that there could be two sub-populations of cells with distinct loss kinetics. Indeed, upon visual inspection of the cells, we noticed that a fraction of the cells contained a large aggregate localized to the old pole (i.e. the pole not renewed after cell division), while the rest contained many small and dynamic aggregates (Fig. 1f-g). This old-pole aggregate was mostly immobile, presumably because its size sterically prevents diffusion through the nucleoid (Fig. 1g). These cells contained bona fide prion aggregates as their progeny contained small aggregates similar to the small and dynamic aggregates that we observed for the other cells in the device (Fig. 1g). We thus re-analyzed the loss kinetics, but this time separately for the small and old-pole aggregate types. We used two different methods for classifying old-pole aggregates, based on the mobility of the aggregates or the fluorescence intensity, which gave similar results (SI Materials and Methods 2.5.3.3.3, Fig. S4a-b). We found that the small aggregates were lost relatively quickly, while the old-pole aggregates were generally much more stable (Fig. 1f-g). The loss curve for the small aggregates fitted well with an exponential decay with a half-life of ~1.5h (Fig. 1f), representing a process with a constant probability of losing the prion state over time (i.e. a memoryless process). This memoryless process is consistent with previous replating experiments, where a similar fraction of prion-positive colonies is found upon successive replating (28, 29, 33). In contrast, few cells with the old-pole aggregates lost the prion over the course of the experiment (93 out of 790 cells), which precluded us from analyzing the loss dynamics of the old-pole aggregates.

These data suggest two modes of prion propagation in *E. coli*: cells containing highly stable old-pole aggregates that give rise to a small aggregate-containing daughter cell at division, and small aggregate-containing cells that lose their prion aggregates with exponential decay. The old-pole aggregate-containing cells would represent a very small fraction of a growing culture (e.g. after 10 divisions, one old-pole cell would become 1 out of 2^10^ = 1,024 cells), but they are enriched in our microfluidic device as we are tracking the cells at the end of dead-end trenches. We thus focused the following analyses on the cells containing small aggregates only.

### Prion loss is mainly driven by partitioning errors at cell division

How do cells lose the prion? A previous study in *E. coli* cells producing the yeast Sup35 PrD suggested that loss of the Sup35 prion could occur through fluctuations in the concentration of the prion protein (33). Based on previous studies in bacteria and yeast (33, 35), we hypothesized that the loss could be due to two non-mutually exclusive mechanisms: 1) stochastic variation in the concentration of the prion protein (or other cellular components, such as the disaggregase ClpB, which is required for the propagation of the *Ch* SSB prion), or 2) mis-partitioning of prion aggregates at cell division (Fig. 2a). These hypotheses lead to different predictions about the prion loss dynamics. If prion loss is caused by stochastic fluctuations in either the prion protein or cellular components, prion loss would be uncorrelated with cell division. On the other hand, if prion loss is caused by asymmetric partitioning of aggregates, the loss would be correlated with cell division and would occur in only one of the two daughter cells.

**Fig. 2.**
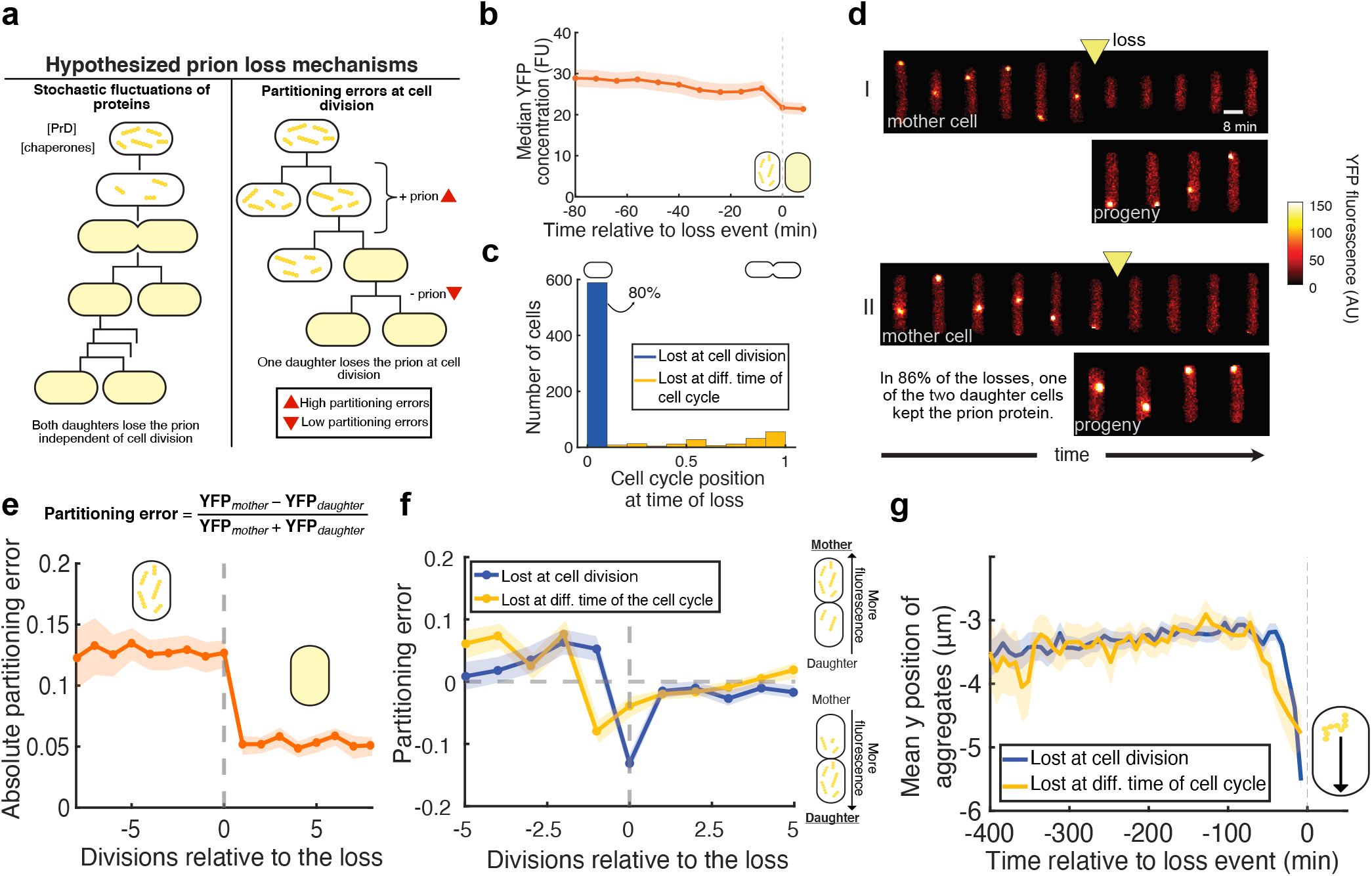
Prion loss is driven by partitioning errors at cell division. **a)** Schematic representation of the hypothesized mechanisms for prion loss in bacterial cells. **b)** Median concentration of fluorescence (Ch SSB PrD) relative to the loss of the prion is constant (*n* = 762 cells). The loss event is indicated with a dotted gray line at time 0. **c)** Histogram of the cell cycle position at the time of loss, where 0 is defined as the moment right after a division and 1 right before. Most cells (~80%) lose the prion immediately after cell division (*n* = 762 cells). **d)** Kymographs of loss event show that prion loss happens in only one of the two daughter cells (86% of the losses, *n* = 356 loss events). YFP fluorescence is shown false colored according to the colormap indicated on the graph. **e)** Mean absolute partitioning errors at the cell divisions relative to prion loss (*n* = 349 cells). The absolute partitioning error is constant prior to the loss, and higher than after the loss. **f)** Mean partitioning errors in the cell divisions relative to the loss show that fluorescence is being transmitted to the daughter at the moment of loss for cells that lost the prion at the moment of cell division (blue lines, *n* = 349 cells). For the cells that lost the prion at a different moment of the cell cycle, this transfer happens one division prior to the loss (yellow line). For symmetric divisions, the average partitioning error would be ~0, since molecules have an equal chance of being inherited by the mother or daughter cells. **g)** Average longitudinal position (y) of tracked aggregates shows that they move toward the daughter cell prior to the loss (*n* = 754 cells). The envelopes represent 2xSEM in **b** and **e-g**.

By tracking prion loss in hundreds of cells with fluorescence microscopy, we could test these hypotheses. Aligning the cells at their moment of loss showed that the fluorescence was constant prior to the loss (Fig. 2b), suggesting that fluctuations in prion protein levels likely play only a minor role in the overall loss. To investigate the possibility that variation in cellular components plays a role in the loss of the prion, we measured the position in the cell cycle at the moment of loss. We observed that ~ 80% of cells lost the prion at the first time point after cell division (Fig. 2c). We also observed that in ~ 86% of losses in the mother (the cell tracked for the duration of the experiment), the prion was maintained in the newly born daughter cell (SI Materials and Methods 2.5.3.4, Fig. 2d). These observations suggested that prion loss is mainly caused by partitioning errors at cell division rather than fluctuations of cellular components, although they do not exclude the possibility that such fluctuations could contribute (36).

Although *E. coli* divides symmetrically with proteins randomly partitioned in the daughter cells, one cell can end up with more of a particular protein by chance. These “partitioning errors” – defined as the normalized difference in the number of molecules between the daughter cells at division (Fig. 2e) – follow a binomial distribution and are generally low because on average they are inversely proportional to the square root of the number of molecules (37, 38). However, cells with the prion have relatively large aggregates, effectively reducing the number of molecules to partition. Partitioning errors at cell division were indeed on average larger and there were more frequent extreme errors (i.e. >30%) before cells lost the prion than after (Fig. 2e, Fig. S5). In addition, the partitioning errors were constant prior to the loss (Fig. 2e), suggesting that the distribution of aggregate size was constant prior to the loss, and that this loss is a sudden rather than gradual event. This further supports the concept that the prion is being propagated until a stochastic event causes its loss. For the cells that lost the prion at cell division, proteins were found to be asymmetrically separated to the daughter at the moment of loss (Fig. 2f). Here, we again define the “mother” cell as the cell tracked for the duration of the experiments, and the “daughter” cells as the progeny that are eventually washed out from the device. For the cells that lost the prion at a different time during the cell cycle, a similar mis-partitioning into the daughter cell was observed one division prior to the loss (Fig. 2f), suggesting that partitioning errors also play a role in the loss of the prion in these cells. Corroborating these results, tracking the position of visible aggregates revealed that they moved on average one cell length towards the daughter cell prior to both types of loss (Fig. 2g). We thus concluded that, at least in this system, prion loss is mainly caused by stochastic partitioning errors of aggregates at cell division, prior to or at the moment of loss.

### Orthologous cPrDs can form prions with similar properties

The two modes of propagation and the molecular events leading to the prion loss could be specific to the studied *Ch* SSB PrD or a more general property of bacterial prions. To begin to investigate this question, we constructed fluorescent fusions of cPrDs from SSB orthologs. We discovered two orthologous SSB PrDs – from *Lactobacillus heilongjiangensis* (*Lh*) and *Moraxella lincolnii* (*Ml*) – that could form self-propagating aggregates after transient expression of the initiation factor New1, as shown with fluorescence microscopy, SDD-AGE, and replating experiments (Fig. 3a-c, S6c). We then evaluated the properties of the aggregates formed by these PrDs in our microfluidic device. Remarkably, we found that their modes of propagation (i.e. small vs old-pole aggregates, Fig. S6a), loss kinetics (Fig. 3a), fraction of loss at cell division (Fig. 3d), and partitioning errors (Fig. S6b) were similar to those formed by the *Ch* SSB PrD (though with some quantitative differences in average loss rates). Therefore, these results support the idea that the modes of prion propagation and the mechanism of prion loss through mis-partitioning at cell division are not only specific to *Ch* SSB PrD, but a more general characteristic among SSB PrDs.

**Fig. 3.**
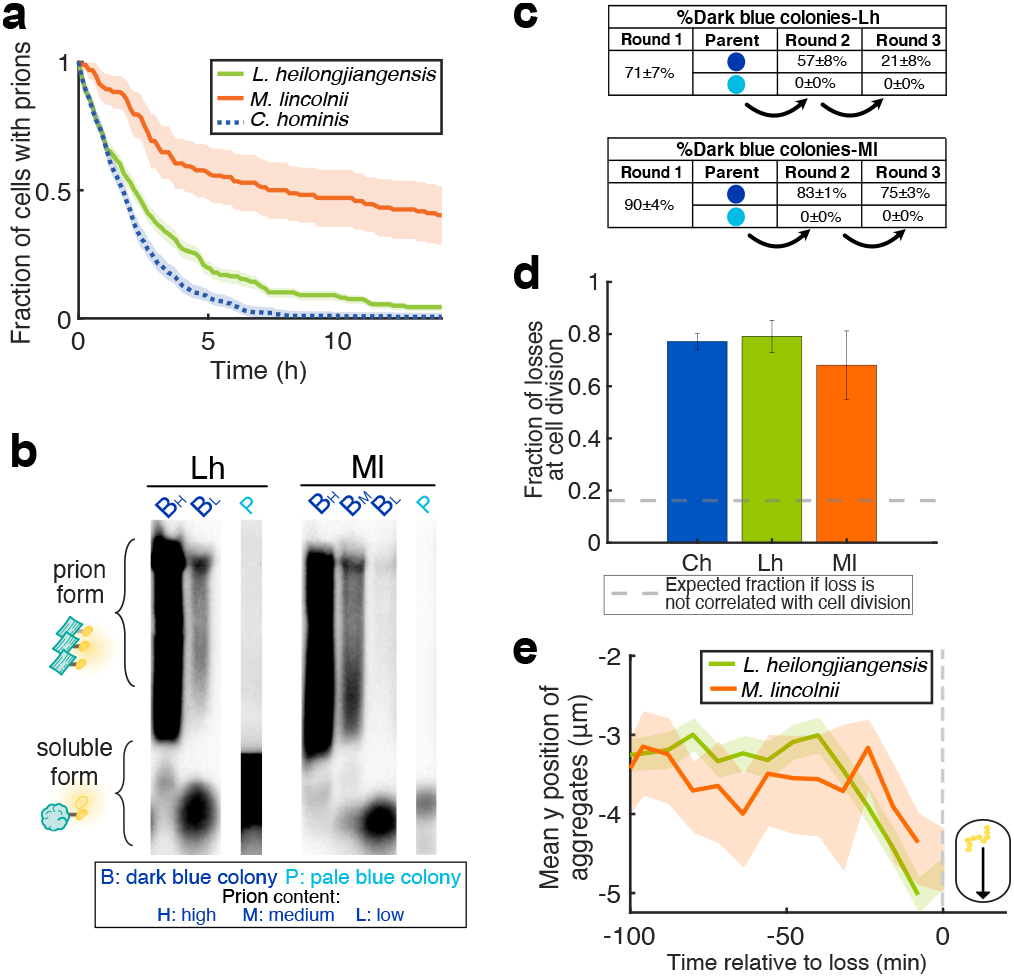
Orthologous SSB cPrDs form self-propagating aggregates comparable to *Ch* SSB. **a)** Prion loss curve for small aggregate cells of *Lh* SSB PrD (*n* = 228 cells) and *Ml* SSB PrD (*n* = 83 cells) compared to *Ch* SSB PrD from Fig. 1. **b)** SDD-AGE of dark and pale colonies confirms the presence of the aggregated prion form of *Lh* SSB and *Ml* SSB in cell extracts derived from dark blue colony cultures. Dark blue colonies with high, medium and low prion content as estimated from fluorescence microscopy images were assayed (Fig. S6d, SI 3.1). Pale colony cultures give rise exclusively to the soluble form. **c)** SSB orthologs form self-propagating aggregates for multiple generations. Replating dark blue colonies gives a mix of dark and pale colonies, while replating pale colonies results in exclusively pale colonies. **d)** Fraction of prion losses at cell division shows that most loss happens at cell division for the different orthologs (*n* = 754 cells for *Ch*, 187 cells for Lh, 47 cells for *Ml*). The error bars represent 2xSEM as estimated by bootstrapping. **e)** Average longitudinal position (y) of tracked aggregates shows that they move toward the daughter cell prior to the loss for the different orthologs (*n* = 187 cells for Lh, 47 cells for Ml). The envelopes represent 2xSEM in **a** and **d-e**.

### A PrD can be propagated with distinct kinetics in distinct lineages

To investigate whether or not these PrDs could form phenotypically distinguishable prion strains (19–21), we quantified prion stability in cells derived from different dark blue colonies representing different lineages propagating the prion. Our experimental setup provided precise and reproducible measurement of the stability; cells containing the *Lh* SSB PrD in its prion form (i.e. the *Lh* SSB prion) and obtained from one colony exhibited similar loss kinetics during four different experiments on four different days (Fig. 4a). However, during our quantification of loss kinetics, we discovered one lineage of cells containing the *Ch* SSB prion that exhibited unusually stable propagation. Quantifying prion stability in cells obtained from this colony in our microfluidic device revealed a loss rate an order of magnitude lower than that of the other lineages (Fig. 4b). To test whether this property was self-propagating, we grew the lineage used for the microfluidic experiments for two additional rounds of about 37 generations each, loading cells from each of the successive rounds of growth into the device (Fig. 4b) and also replating them on indicator medium (Fig. S7a). Strikingly, the loss kinetics were constant over ~110 generations and nearly all colonies were prion positive after each round of plating. DNA sequencing of the PrD-containing plasmid from cells of this lineage revealed no mutation in the promoter, the PrD, or the plasmid origin of replication (Fig. S7d), suggesting that the stability property is inherited through the structure of the aggregates rather than genetically.

**Fig. 4.**
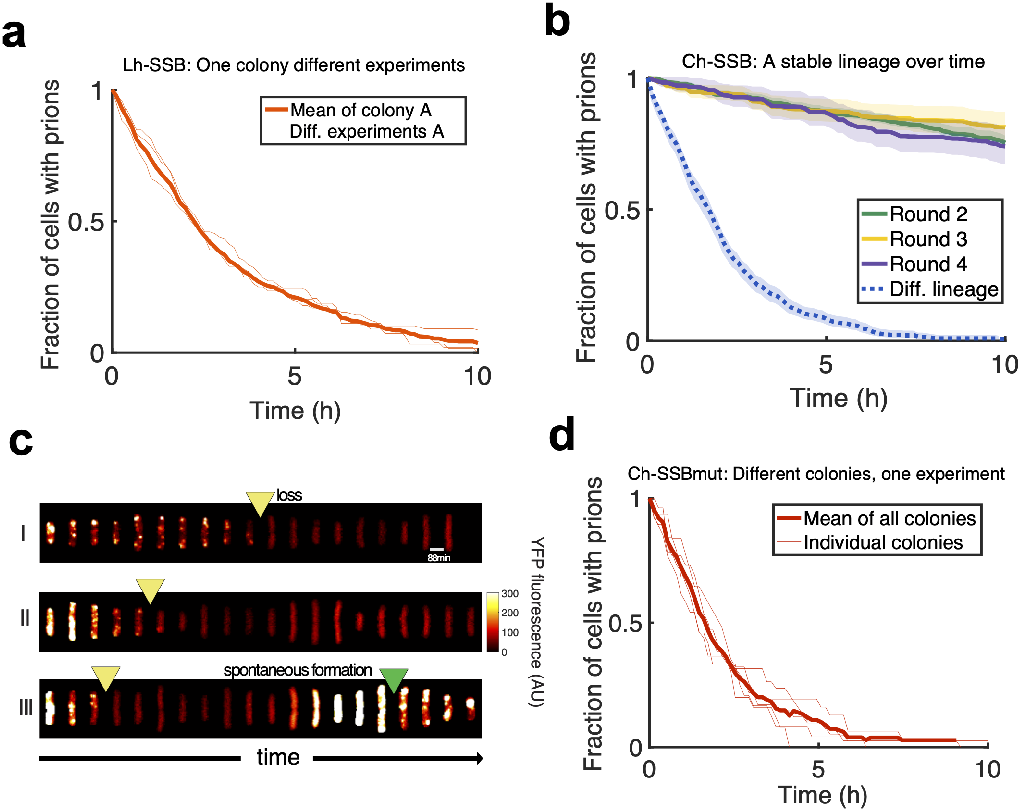
Distinct bacterial lineages propagating identical prion protein exhibit distinct prion loss kinetics. **a)** The experimental setup provides precise measurement of the prion loss kinetics. Prion loss curves for one colony of *Lh* SSB PrD in four different experiments (thin orange lines, average in bold, *n* = 815 cells total). **b)** The prion loss curve for a stable lineage of *Ch* SSB PrD remains constant over multiple rounds of growth (~37 generations each, *n* = 1,018 cells total). Round 1 refers to the first plating of induced cells cured of New1, and each subsequent round includes an overnight growth in liquid culture and plating on indicator medium. Round 2, 3, and 4 cells were obtained from a colony culture inoculated from a Round 2, 3, and 4 colony, respectively. Another lineage (from Fig. 1, dashed blue line)) is shown as a comparison. The envelopes represent 2xSEM as estimated by bootstrapping. **c)** Kymograph of a mutant of *Ch* SSB PrD that can form self-propagating aggregates without the presence of the initiation factor (termed *Ch* SSBmut PrD). YFP fluorescence is shown false colored according to the colormap indicated on the graph. The prion is eventually lost, but rare spontaneous re-formation (green arrow) happens at low inducer concentration (2 *μ*M IPTG for the duration of experiment). The spontaneous re-formation events were observed following large stochastic fluctuations in fluorescence, likely due to plasmid copy number variation. Such fluctuations were also observed in experiments with other PrDs, but in these cases they did not cause re-formation of the prion. **d)** Prion loss curve for different colonies of the *Ch* SSBmut PrD exhibit similar propagation dynamics (thin line, average in bold, *n* = 155 cells).

### A mutant PrD can form a prion without an initiation factor

To explore the possibility that genetic mutations can be identified that increase prion-forming propensity, we performed random mutagenesis of the PrD-encoding moiety of the *Ch* SSB PrD construct (SI Materials and Methods 2.1.2). We screened for and isolated a mutant (termed *Ch* SSBmut PrD) that formed dark blue colonies with SDS-stable aggregates even without exposure to the initiation factor New1 (Fig. S8b). To investigate whether or not the aggregates formed by this mutant were propagated in a similar manner to those formed by the *Ch* SSB PrD, we characterized the dynamic properties of the *Ch* SSBmut PrD in the microfluidic device. We observed that the mutant aggregates were propagated and lost with similar modes, kinetics, and loss mechanisms as the wild-type *Ch* SSB aggregates (Figs. 4c-d, S8a-f). However, in some rare cases, the mutant protein could spontaneously reform the aggregates, consistent with its ability to access the prion conformation independently of New1 (Fig. 4c). We speculate that the *Ch* SSBmut PrD is a prion domain with a high probability of forming one particular strain. Consistent with this possibility, we found that cells from five distinct colonies exhibited the same kinetics of prion loss (Fig. 4d). This mutant will be characterized extensively in another study. Nonetheless, our results indicate that the aggregates formed by the *Ch* SSBmut PrD are bona fide prions despite their [*PIN*^+^]-independence.

### Physiological impact of the presence of prions aggregates

We then sought to determine the general physiological impact of such heterologous prion aggregates in *E. coli*. Among eukaryotic prions, it is striking that some are the cause of fatal neurodegenerative diseases while others appear to have low or no toxicity (1, 9–11, 17). In bacteria, a potential impact on growth rate (as a proxy for cell viability) is challenging to precisely quantify in bulk due to the different modes of prion propagation as well as the stochastic loss of the prion during growth. Thus, using our microfluidic device, we quantified the growth rate of individual cells that did not have the prion, of cells that maintained old-pole aggregates, and of the cells with small aggregates. Cells with small aggregates had a median doubling time ~1.5% slower than cells without the prion, and cells with old-pole aggregates had a ~3% growth penalty compared to cells without the prion (Fig. S9a). We also quantified the death rate of cells propagating the prion, and observed that the death rate was overall very low (~6×10^−3^ /h) and similar to cells not propagating the prion (Fig. S9b). We thus concluded that the presence of prion aggregates had a small yet meaningful negative effect on the overall cell physiology.

### A stochastic model recapitulates the experimentally observed prion propagation dynamics

Prion propagation in yeast and mammals has been mathematically modeled in various studies (19, 22, 23, 25, 39–41). To investigate if these molecular models can describe the observed dynamics of our system, we adapted a mathematical model of prion propagation for single bacterial cells. In particular, we modeled the propagation and loss of prion aggregates in growing and dividing cells with a stochastic generalization of the nucleated polymerization model (24, 25) (Fig. 5a, details in SI 3.2). Proteins are produced in the soluble form, and can then be converted in the prion form by elongation of an existing aggregate oligomer. Aggregates can be fragmented into smaller oligomers – keeping the number of monomers constant – and aggregates below a critical size *n* spontaneously fold back into the soluble form. Cells grow continuously and divide once they reach a critical size, such that proteins are randomly partitioned between the two daughter cells according to a binomial distribution (37, 38). Individual time traces were generated using the Gillespie algorithm, which simulates the stochastic chemical reactions (42).

**Fig. 5.**
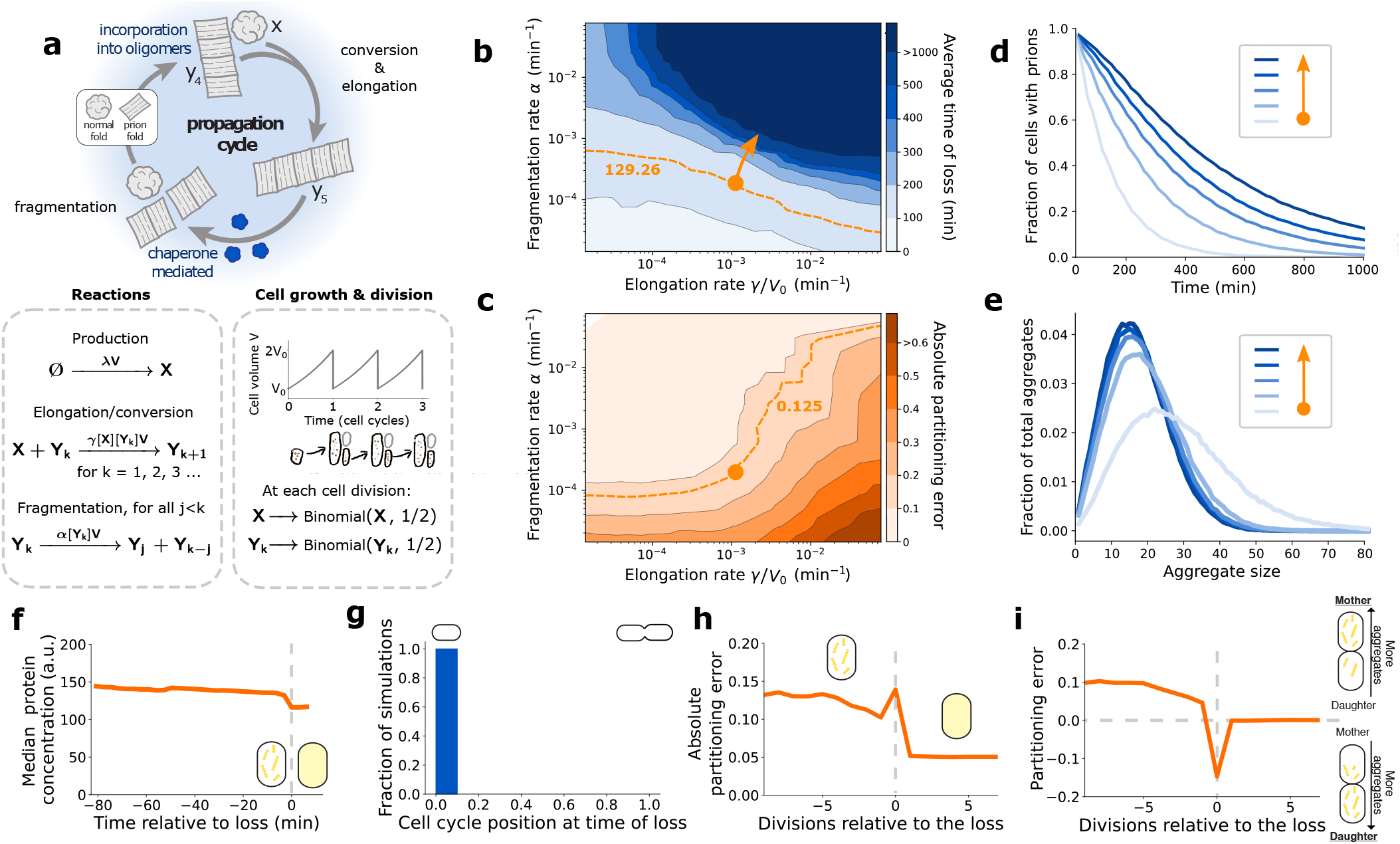
A stochastic nucleated polymerization model recapitulates the experimental results. **a)** A stochastic model of prion propagation in growing and dividing cells. Soluble fold protein numbers, denoted by X, are produced constitutively with a rate that scales with the cell volume, so that their concentration becomes cell-cycle independent (see SI 3.2.1). The number of prion fold aggregates made of *k* proteins is denoted by *Y_k_*, where *k* = 1, 2, 3,…. When a soluble fold protein collides with an aggregate of size *k*, it can be converted to prion fold by elongating the aggregate to size *k* + 1. Assuming mass action kinetics, soluble fold proteins are converted to prion fold with a reaction rate proportional to the protein concentrations. Similarly, chaperon mediated fragmentation follows a reaction that is proportional to the aggregate concentrations, with each binding between any two monomers having the same probability of splitting. Concentrations are given by dividing the protein numbers by the cell volume, which grows exponentially from *V*_0_ to 2*V*_0_ between divisions with a fixed doubling time. At cell division, protein numbers are split randomly, with each soluble protein and each aggregate having a 50% chance of remaining in the cell. **b)** Soluble fold production parameter *λV*_0_ was estimated to be 1.75 min^−1^ by comparing the measured partitioning error of cells after loss of prions with their respective simulations (see SI 3.2.3.2). With no minimal seed size *n* = 0 (see SI 3.2.5 for *n* = 2), a parameter sweep of elongation and fragmentation parameters shows that prions in cells with larger fragmentation and elongation rates are more stable. An average time of loss of 129.26 min was measured in the experiment shown in Fig. 1f, with the corresponding contour indicated by the dashed orange line. **c)** Cells with smaller fragmentation rates and larger elongation rates have larger partitioning error prior to loss. An absolute partitioning error prior to loss of 0.125 was measured in the experiment shown in Fig. 2e, with the corresponding contour indicated by the dashed orange line. Using the two contour plots from **b** and **c** we find the model parameters that match the measured time of loss and partitioning error, indicated by the orange dot. **d)** Time of loss curves follow an exponential, in agreement with Fig. 1f. Plotted are the time of loss curves for systems with parameters along the solid orange line in **b**. Loss is defined as when *Y_k_* = 0 for all *k*. **e)** The model can predict the aggregate size distribution prior to loss, showing that smaller aggregates are more stable in this parameter regime. **f)** The total protein concentration is approximately constant leading up to the loss, in agreement with Fig. 2b. **g)** In this model the prion state is always lost at cell division. **h)** Absolute partitioning errors are larger before the loss, in agreement with Fig. 2e. **i)** A large negative partitioning error occurs at the time of loss, in agreement with Fig. 2f.

First, we simulated the model in a large parameter space of elongation and fragmentation rates (Fig. 5b-c). We found that systems with large elongation and fragmentation rates were more stable as they take longer to lose the prions. Outside of this parameter space, however, the prion was eventually lost on timescales similar to our experiments. We then estimated the elongation and fragmentation rates by selecting the unique model parameters that matched the observed loss rates and partitioning errors as indicated in Fig. 5b-c (see SI 3.2.3 for details). Strikingly, this simple model could recapitulate all the observations from the experiments. We find that simulated cells reached a quasi-stationary state, where the distribution of prion aggregates (Fig. S10a-d), the total amount of protein (Fig. 5f), and the absolute size of partitioning errors (Fig. 5h), were approximately constant prior to the loss. As observed experimentally, a large partitioning error into the untracked cell was observed at the moment of loss (Fig. 5i), which happened at cell division (Fig. 5g). Finally, the loss curve in the population followed an exponential decay, corresponding to constant probability of loss over time (Fig. 5d). The model also shows how different prion conformations, with potentially different elongation and fragmentation rates, can lead to different stabilities.

Using this model, we predicted that cells with larger volumes would have lower partitioning errors, which would make the prion more stable (Fig. S10e-f). To test this prediction, we used a mutant with longer cell size but with the same growth rate (*ftsN* deleted of codons encoding amino acid residues 244-319, (43), Fig. S11a,b,d), which revealed that ~ 50% fewer cells lost the prion during replating experiments (Fig. S11c). We thus concluded that partitioning errors played an important role in the loss of the prion, that cell volume affects prion loss, and that the nucleated polymerization model was consistent with our experimental results.

## Discussion

Here, we used microfluidics and fluorescence microscopy to track thousands of individual cells propagating prion aggregates. Notably, cells tracked for over 20 generations with the prion would have likely renewed almost every single protein in the cell (and thus the prion proteins many times), showcasing the self-propagating nature of the prion aggregates. For proteins that are not degraded, half of the proteins are renewed after one cell division. Thus, after 20 cell divisions, 1/2^20^ of the ~ 2^21^ original proteins will not have been renewed, such that only a handful of the original proteins will remain (44, 45).

### Modes of propagation

We discovered that, for the three PrDs studied, the prions were propagated through two modes: stable old pole aggregates and less stable small aggregates (Fig. 6).

**Fig. 6.**
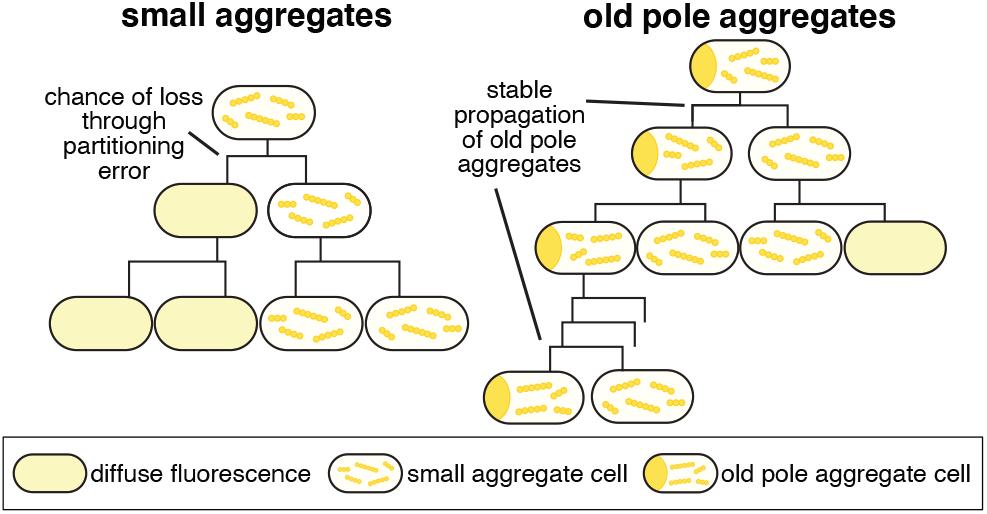
Schematic of the two observed modes of prion propagation. Cells with small aggregates have a probability of losing the prion at each cell division through partitioning errors. At cell division, an old-pole aggregate cell generates a small aggregate cell and an old-pole aggregate cell. Although the old-pole aggregate is very stable, the cells containing old-pole aggregates represent a small fraction of a growing culture. The small aggregate cells generated through this division presumably propagate the prion similarly to the other observed small aggregate cells.

We note that the old-pole aggregate cells also contain small aggregates which can be difficult to visualize due to the brightness of the large aggregate. Therefore, at division, the old-pole cells generate one cell bearing an old-pole aggregate and one bearing small aggregates (Fig. 6). We have not investigated the formation of these old-pole aggregates, but speculate that they can be formed stochastically once an aggregate reaches a critical size. This critical size would prevent them from freely diffusing through the cell and confine them to the pole, while potentially also preventing chaperones from fragmenting them normally. It remains to be determined if other PrDs, from bacteria or other organisms, exhibit this type of propagation. Yet, we conjecture that these old-pole aggregate cells could form a rare yet stable reservoir of the prion epigenetic state, generating cells containing small aggregates at each cell division.

In contrast, the cells containing the small aggregates lost the prion relatively quickly, with a constant rate of loss over time (memoryless process with half-life of ~2-6 generations). We note that this stability will depend on the concentration of the prion protein, which was kept as low as possible during these experiments. The loss of the prion in these cells was driven mainly by a sudden mis-partitioning of prion aggregates at cell division, giving a probability of losing the prion at each cell division (Fig. 6), consistent with the memoryless loss kinetics. It remains to be determined if other bacterial PrDs, such as the Rho PrD from *Clostridium botulinum* (which had a lower rate of loss during replating (28)), are propagated and lost similarly.

### Different lineages have different stability

In addition to disentangling the modes of propagation at the single-cell level, our microfluidic assay enabled precise quantification of the loss kinetics. This enabled us to observe that distinct lineages of the same PrD could propagate aggregates with distinct stabilities. In particular, we characterized one lineage of the Ch SSB prion that had a stability an order of magnitude greater than the others. This finding recapitulates and extends observations made in the previous study of the *Ch* SSB PrD, where both low-propagation and high-propagation lineages were characterized (29). These results are reminiscent of what has been observed in yeast, where one protein (e.g. Sup35) can form multiple self-propagating structures, called strains, with different stabilities (e.g. [*PSI^+^*]^strong^ vs. [*PSI^+^*]^weak^) (19, 46–49). Further work will be necessary to show whether the different lineages observed reflect different self-propagating structures.

In contrast, we characterized a mutant PrD that could form self-propagating aggregates without an initiation factor (independently of [*PIN*^+^]). We conjecture that this mutant is a PrD with a high probability of forming one particular self-propagating structure, similar to how certain mutations of the mammalian PrP lead to the formation of a particular prion strain in genetic prion diseases (e.g. familial CJD) (50–52).

### Molecular model of prion propagation and challenges in bacteria

Finally, we developed a stochastic implementation of the nucleated polymerization model that could recapitulate all the observed single-cell properties. In the future, the simple model could be tested further by perturbing the experimental parameters, e.g. by changing the concentration of the disaggregase ClpB (required for the propagation of the *Ch* SSB prion). This would indicate whether additional constraints that have been necessary to explain results in yeast, such as a size-dependent transmission of aggregates (41) or different seed size for prion strains (23), are also necessary. This model also reveals challenges for prion propagation in bacteria. Using the experimental measurements (partitioning errors and the average time of prion loss), we can estimate the total number of proteins, the fragmentation rate, and the elongation rate, and thus obtain an approximation for the replication rate (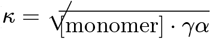, see SI 3.2.6). Even though the PrDs studied here appear to be lost relatively quickly, the estimated replication rate (~ 10^−5^/s) is of similar order of magnitude to other prions, such as the mammalian PrP in vivo (22).

How does the model explain the discrepancy between the fast replication rate and the prion instability? *E. coli* is small and therefore has low numbers of proteins, which results in high partitioning errors. For example, the total number of proteins is ~100 times smaller in *E. coli* than in *S. cerevisiae*, which would result in partitioning errors ~10 times larger (i.e. 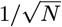). In addition, *E. coli* divides rapidly, which further reduces the stability of the prion, as proteins need to be converted to the prion state prior to the division for stable propagation. The lower stability we observed contrasts with what was observed in yeast, with e.g. a loss rate of 10^−5^ generations^−1^ for [*PSI*^+^]. Nevertheless, we speculate that less stable PrDs do not make them less useful as an epigenetic switch. Prions have been suggested to provide an epigenetic state with fitness advantage under certain environmental conditions (4–6, 15, 16). The optimal stability of such an epigenetic state depends on the rate of change of the environment experienced by the organism, which is difficult to estimate. Thus, whether a loss rate on the order of generations (for the PrDs studied here) or tens of thousands of generations (e.g. yeast [*PSI*^+^]) is more or less useful biologically depends on temporal dynamics of the environment.

In conclusion, this work further establishes the conservation of prion propagation across domains of life. Further work will unravel how many of the thousands of predicted prokaryotic candidate PrDs can form prions, and how prion formation affects cell physiology.

## Materials and Methods

Detailed Materials and Methods are available in the SI Appendix. The base strain used throughout the paper was *E. coli* MG1655. Prion formation was induced overnight by the production of the SSB PrDs fusion proteins and the New1 fusion protein with 10*μ*M IPTG at 30°C. Cells were cured of the New1-containing plasmid by plating overnight at non-permissive temperature (37°C). These indicator plates contained X-Gal which enabled distinguishing colonies with prion-containing cells (dark blue). For the microfluidic experiments, dark blue colonies were grown overnight at 30°C and the cultures were inspected with fluorescence microscopy to confirm that the cells contained prion aggregates. These confirmed cultures were then loaded into the microfluidic device, where the cells were continuously fed a supplemented M9 growth medium. Multiple cell positions were imaged in fluorescence every 8 min with a Zeiss Axio Observer at 63x, and the cell lineages were segmented and tracked as previously done.

### Data and materials availability

The segmented and tracked lineage data will be available on Dryad and the code for analyzing this data and generating the figures in the manuscript will be available on Github. The microscopy time-lapse images are available upon request due to their large size. The plasmids used in this study will be available on Addgene.

## ACKNOWLEDGMENTS

K.J., M.T.O.H., F.P., and G.M. received fellowships from NSERC CREATE SynBioApps fellowship (511601-2018). F.P. and G.M. received Master’s and Doctoral fellowship from NSERC. F.P. received an FRQNT Doctoral fellowship and a Miriam Aaron Roland fellowship. This work was supported by NIH Grant GM136247 (to A.Hochschild), NSERC Discovery grants (RGPIN-2019-07002, to L.P.T. and RGPIN-2019-06443 to A.Hilfinger) and a CFI John R. Evans Leader Fund (38290, to L.P.T.).

