## Supplementary Information for "Measuring prion propagation in single bacteria elucidates mechanism of loss"

**This PDF file includes:**

Figs. S1 to S17
Tables S1 to S2
SI References

### 1. Supplementary Figures

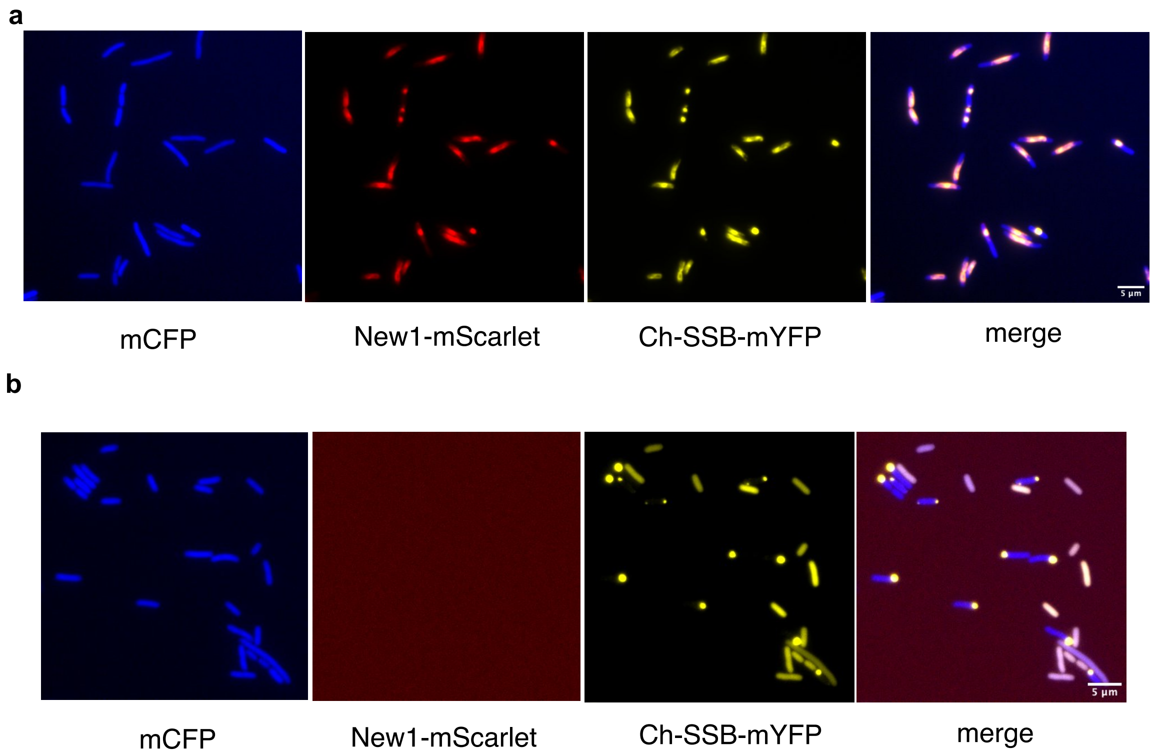

c

| Chloramphenicol | + | - |
|---|---|---|
| CFU/mL | <10 | $1.35 \times 10^{7}$ |
| Frequency of New1 positive cells: $<1 \times 10^{-6}$ | | |

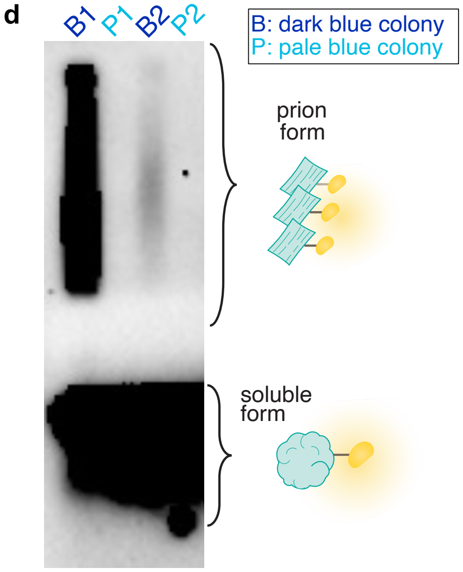

**Fig. S1. Complete curing of New1-mScarlet-I-expressing temperature sensitive plasmid and subsequent induction of prion formation a)** Fluorescence microscopy of cells that harbor *Ch* SSB PrD before curing of the New1-mScarlet-I plasmid. The cells express CFP constitutively (for cell segmentation and tracking), mEYFP (fused to *Ch* SSB PrD) and mScarlet-I (fused to New1). **b)** Fluorescence microscopy of cells that harbor *Ch* SSB PrD after curing the temperature sensitive plasmid at 37°C overnight shows that no New1-mScarlet signal is detected. **c)** Plating of a culture used for microfluidic experiment shows that no cells contain the cured New1-expressing plasmid. CFU/ml of cells grown overnight with and without chloramphenicol (resistance marker of New1-mScarlet-I plasmid). No colonies are observed in the presence of chloramphenicol. **d)** SDD-AGE shows that dark blue colonies contain SDS-stable aggregates, whereas pale colonies contain only soluble *Ch* SSB PrD. Prion formation was previously induced with New1-mScarlet-I and then the cells were cured of the New1-mScarlet-I plasmid.

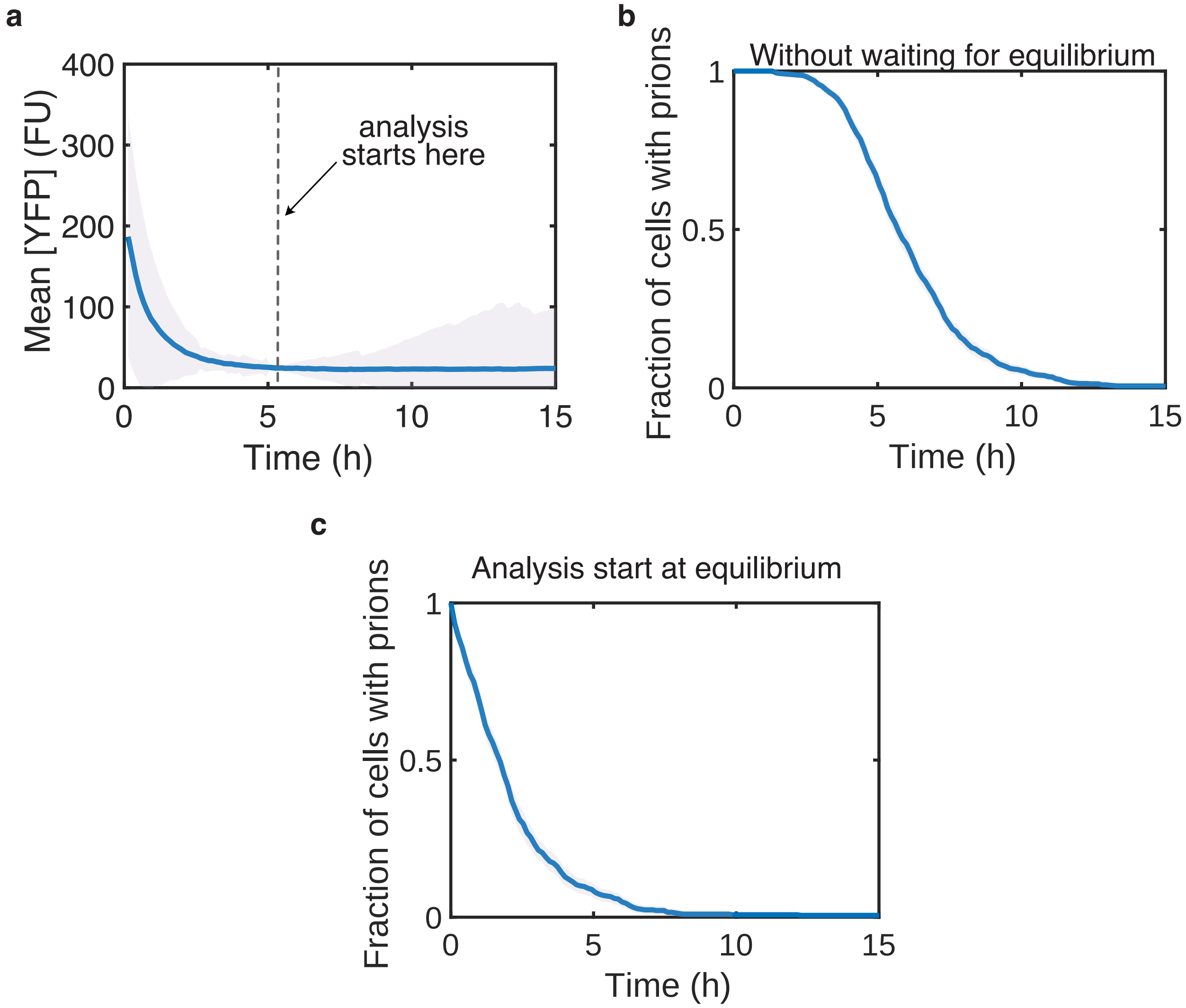

**Fig. S2. Equilibration of growth conditions in the microfluidic device prior to beginning the analysis. a)** The concentration of the *Ch* SSB PrD (as reported by its fused YFP) reaches equilibrium after $\sim$5h in the device. Average concentration of YFP fluorescence for the cells in the device over time (n = 947 cells), where the dashed line indicates where the analysis starts for the experiment. The decrease in YFP fluorescence is due to the change of the concentration of IPTG from 10 $\mu$M outside the device to 0 $\mu$M in the device. **b)** and **c)** Prion loss curve with the analysis starting immediately with the imaging (**b**, n = 947 cells) and with the analysis starting when growth conditions are at equilibrium (**c**, n = 614 cells). The envelopes indicate 2xSEM (estimated by bootstrapping for **b** and **c**).

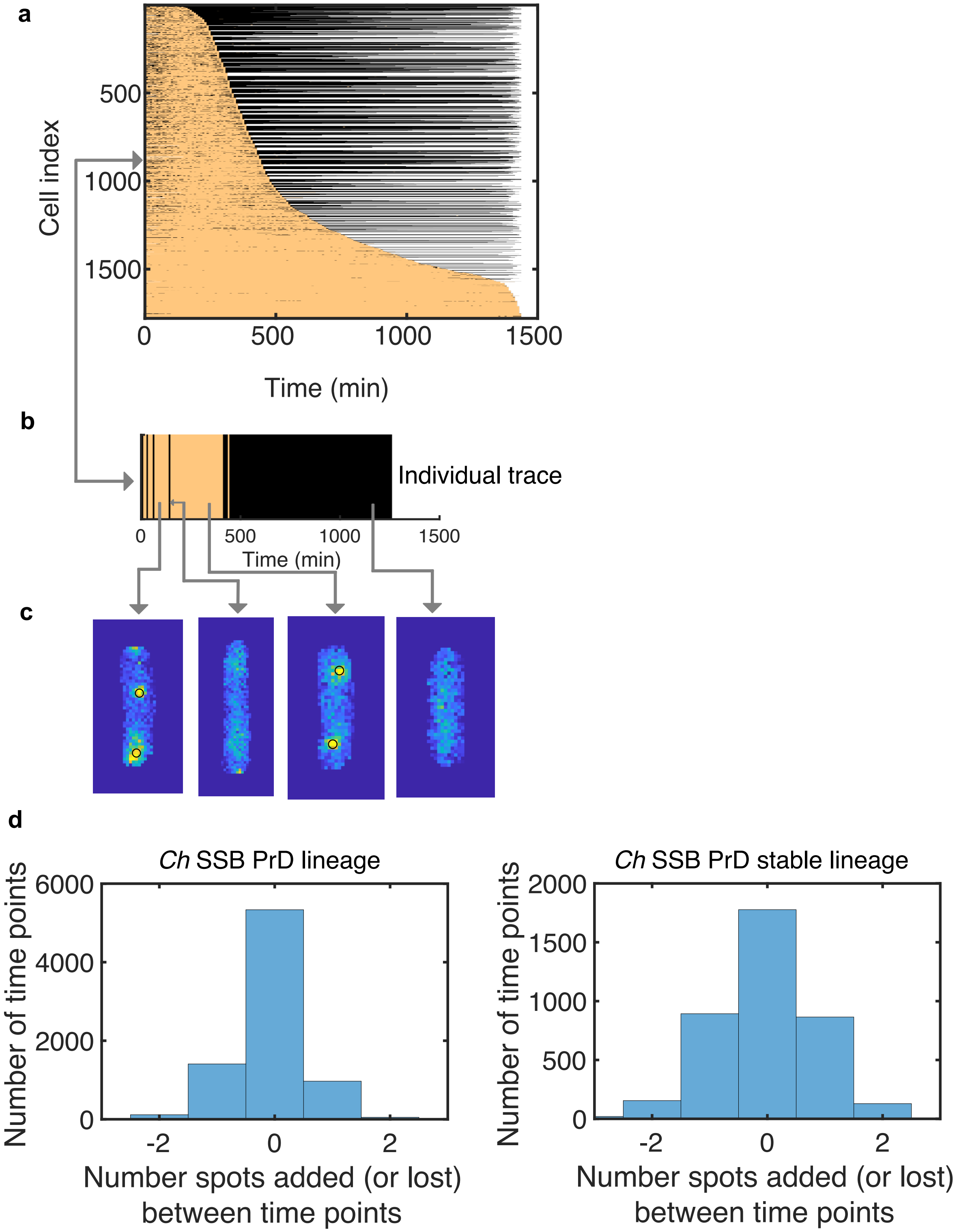

**Fig. S3. Estimation of prion loss using a spot-finding algorithm. a)** Cell traces ordered by the duration of prion propagation, with the coloration indicating the presence (beige) or absence (black) of spots, as determined by a spot-finding algorithm. The prion loss was called the first time 8 subsequent time points have no aggregates. **b)** One example cell trace showing the irreversible loss of the prion aggregates. Aggregates are sometimes not detected during the prion propagation phase, presumably because the size of the aggregates is below the detection limit. **c)** Example images of the cell tracked in **b)**, showing the detection of the aggregates using the spot-finding algorithm (black circles). **d)** Aggregates are added and removed during the tracking of the cells in the device. Histogram of the difference in the number of detected aggregates in between time points for the *Ch* SSB lineage of Fig. 1 (left, n = 586 cells) and the ultra stable lineage of Fig. 4b (right, n = 553 cells).

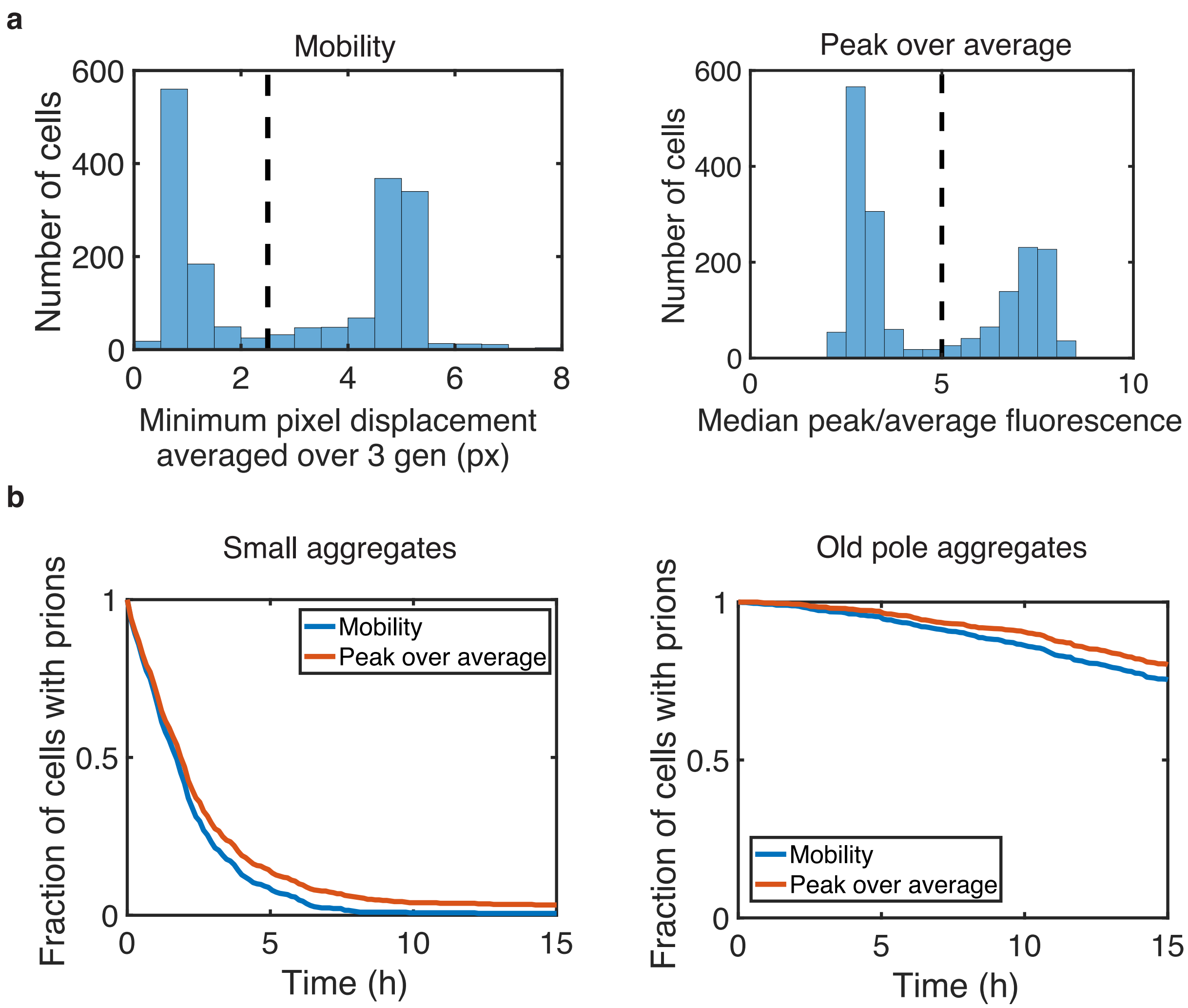

**Fig. S4. Two methods for classifying aggregates give similar results. a)** (Left) Aggregates can be distinguished by measuring movement of the aggregate over time (mobility), with old-pole and small aggregate cells displaying low and high mobility, respectively. Histogram of minimum movement of aggregates averaged over three generations for each cell. (Right) Alternatively, aggregates can be classified by their fluorescence intensity, with old-pole aggregates being brighter. Histogram of peak fluorescence intensity (median of top 10% of pixels in the cells) divided by the average fluorescence intensity. Dashed lines represent the thresholds for classification of aggregates as small or old-pole types (n = 1,788 cells). **b)** Prion loss curve for the small (left, n = 614 cells for mobility and 681 cells for peak over average) and old-pole aggregates (right, n = 831 cells for mobility and 764 cells for peak over average), with the classification performed using the mobility (blue) or the fluorescence intensity (red) thresholds of **a**).

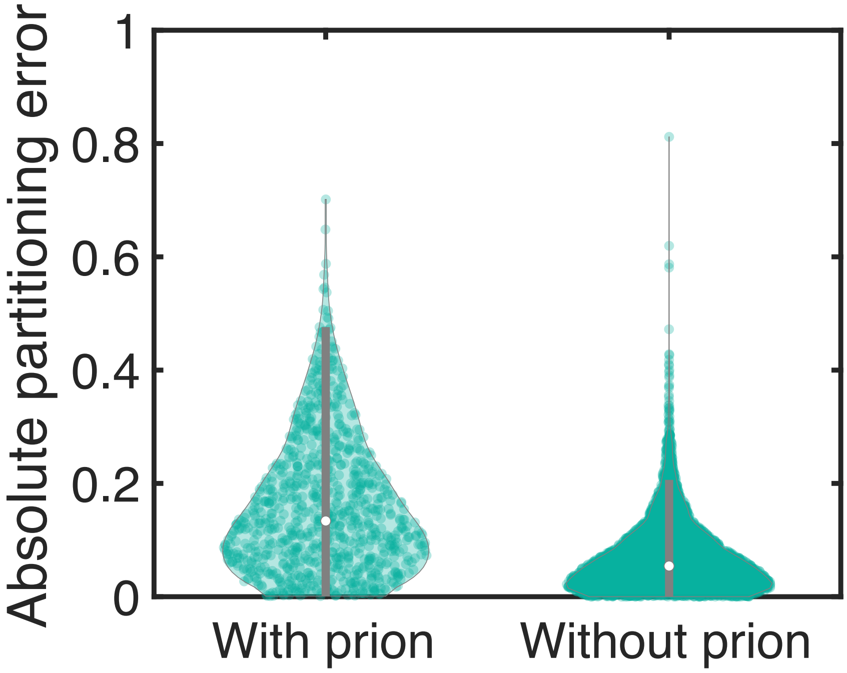

**Fig. S5. The partitioning errors are greater when the cells contain prions.** Violin plot showing the distribution of absolute partitioning errors ($|YFP_{mother} - YFP_{daughter}|/(YFP_{mother} + YFP_{daughter})$ for cells that contain prion and after they lost the prion (n = 886 cell division with the prion, 4,997 divisions without the prion). The partitioning errors are on average larger and there are more frequent extreme partitioning errors when cells contain prion aggregates.

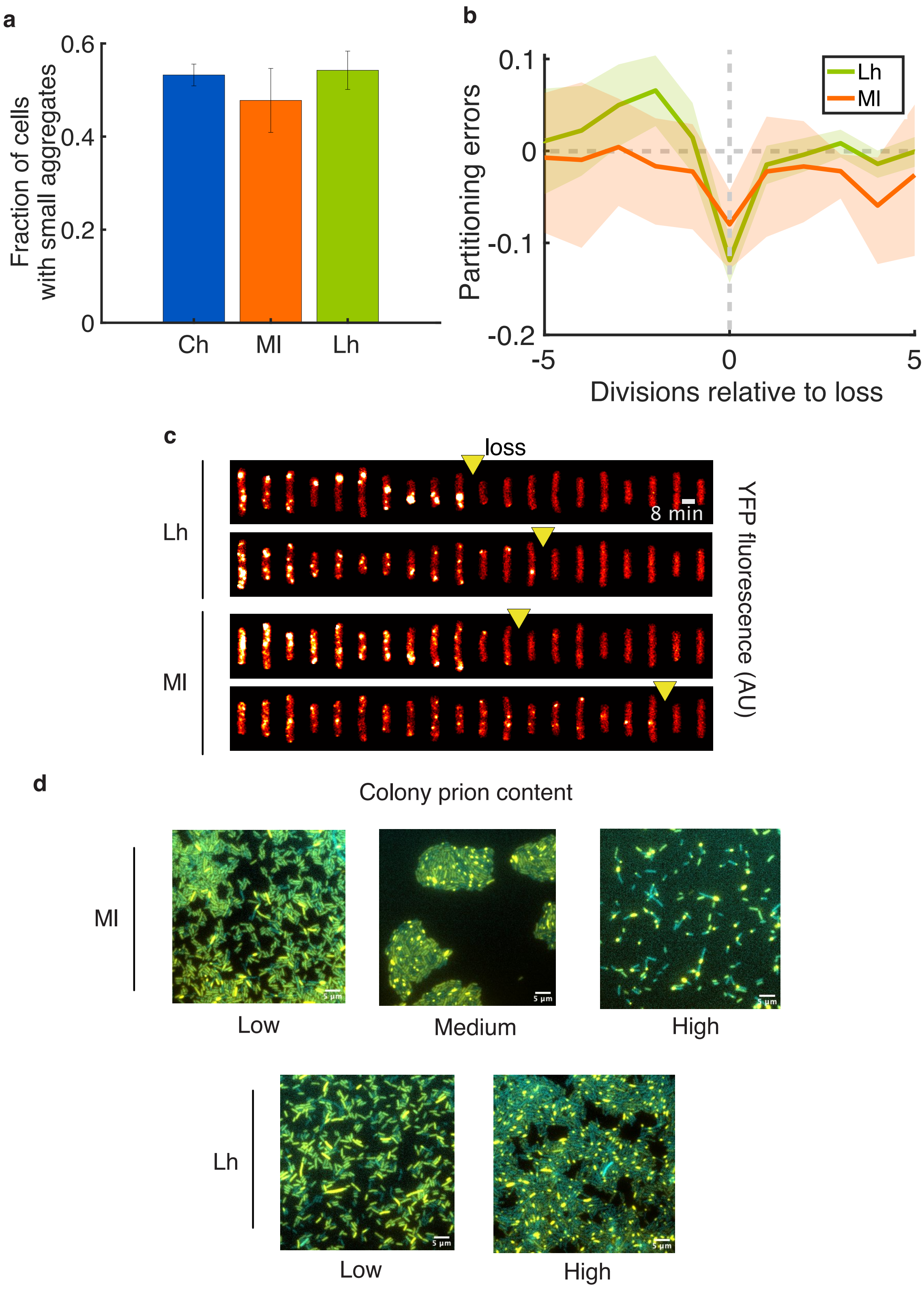

**Fig. S6. Orthologous SSB PrDs form self-propagating prion aggregates with similar properties. a)** Fraction of cells that contain small aggregates (as opposed to old-pole aggregates) for cells containing the *Ch* SSB (blue, n = 1,779 cells), *Ml* SSB (orange, n = 224 cells ), or *Lh* SSB (green, n = 601 cells) prions. **b)** The orthologs also show a transfer of fluorescence to the untracked cell at the moment of prion loss, as shown in the partitioning errors in the divisions relative to the loss (*Lh*, green line, n = 99 cells, *Ml*, orange line, n = 29 cells). **c)** Kymographs showing the propagation and loss of the prion aggregates for the two orthologs, where the yellow triangles indicate the losses identified using the spot-finding algorithm. **d)** Fluorescence microscopy of dark blue colonies with *Lh* SSB PrD and *Ml* SSB PrD. The YFP is shown in yellow (SSB PrD fusion) and the CFP (constitutive marker for segmentation) is shown in cyan. Cells with prions show aggregated fluorescence, and cells without prions show diffuse fluorescence. The fraction of cells with prions can differ between different colonies, presumably due to the stochastic loss of the prion during the colony forming process.

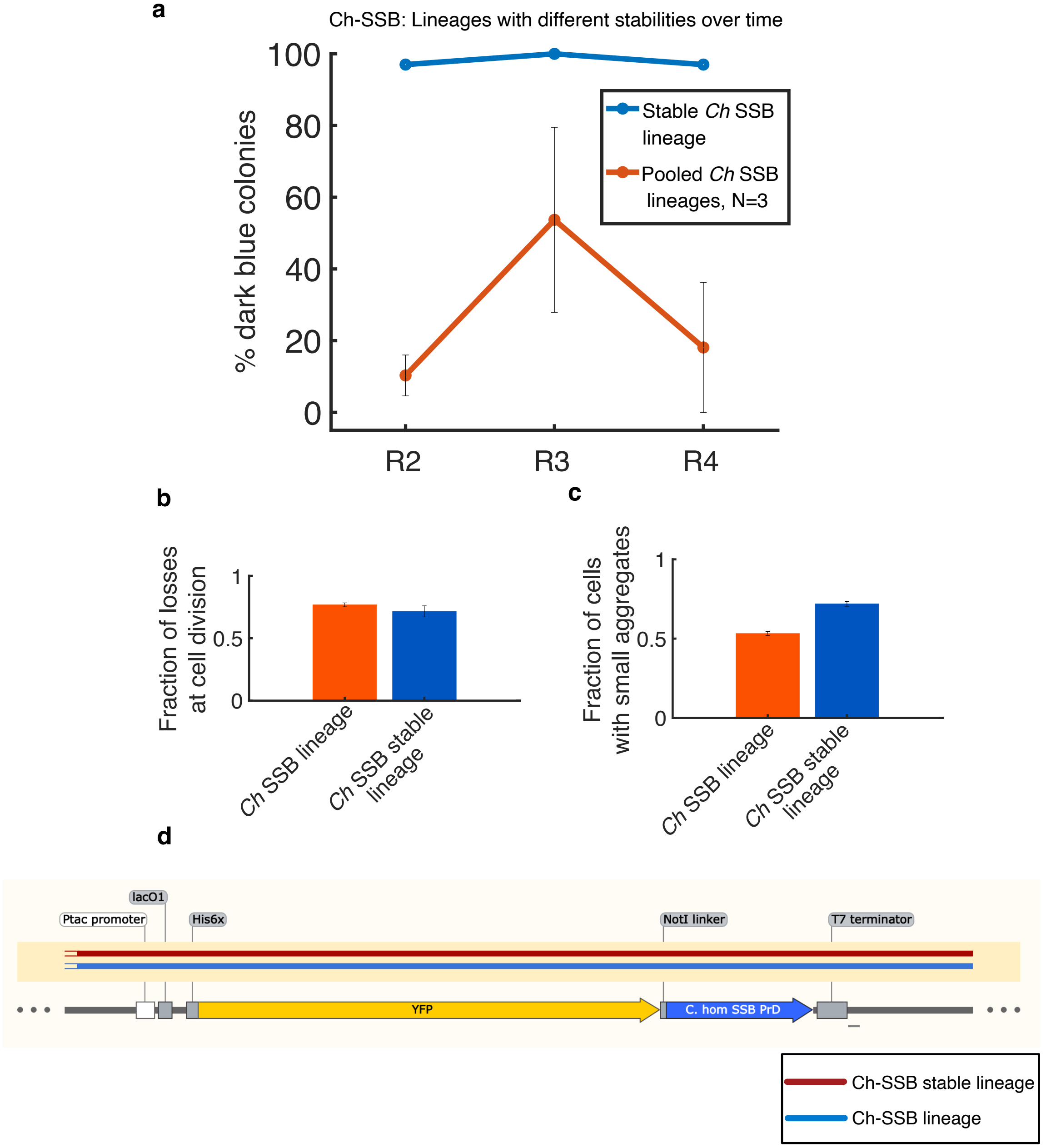

**Fig. S7. One lineage propagates the *Ch* SSB prion with very high stability. a)** The stability of the prion is kept over multiple rounds of growth. The stable lineage (blue) and a pool of *Ch* SSB prion-containing colonies (red) were grown and replated for multiple rounds, and the fraction of colonies with prion-containing cells was estimated using the P*clpB-lacZ* reporter. R2, R3, and R4 designate the colonies obtained after plating the Round 1, Round 2, and Round 3 colony cultures, respectively (see Fig. 4b). While the pooled colonies show a fraction of colonies with prion-containing cells between 10% and 50%, the ultra-stable lineage has close to 100% during each replating. The properties of the ultra-stable lineage are similar in the microfluidic device, where most losses happen at cell division **(b)**, and with a similar fraction of cells with small aggregates **(c)**. **d)** DNA sequencing of the plasmid containing the PrD-YFP fusion in the stable lineage shows no mutations.

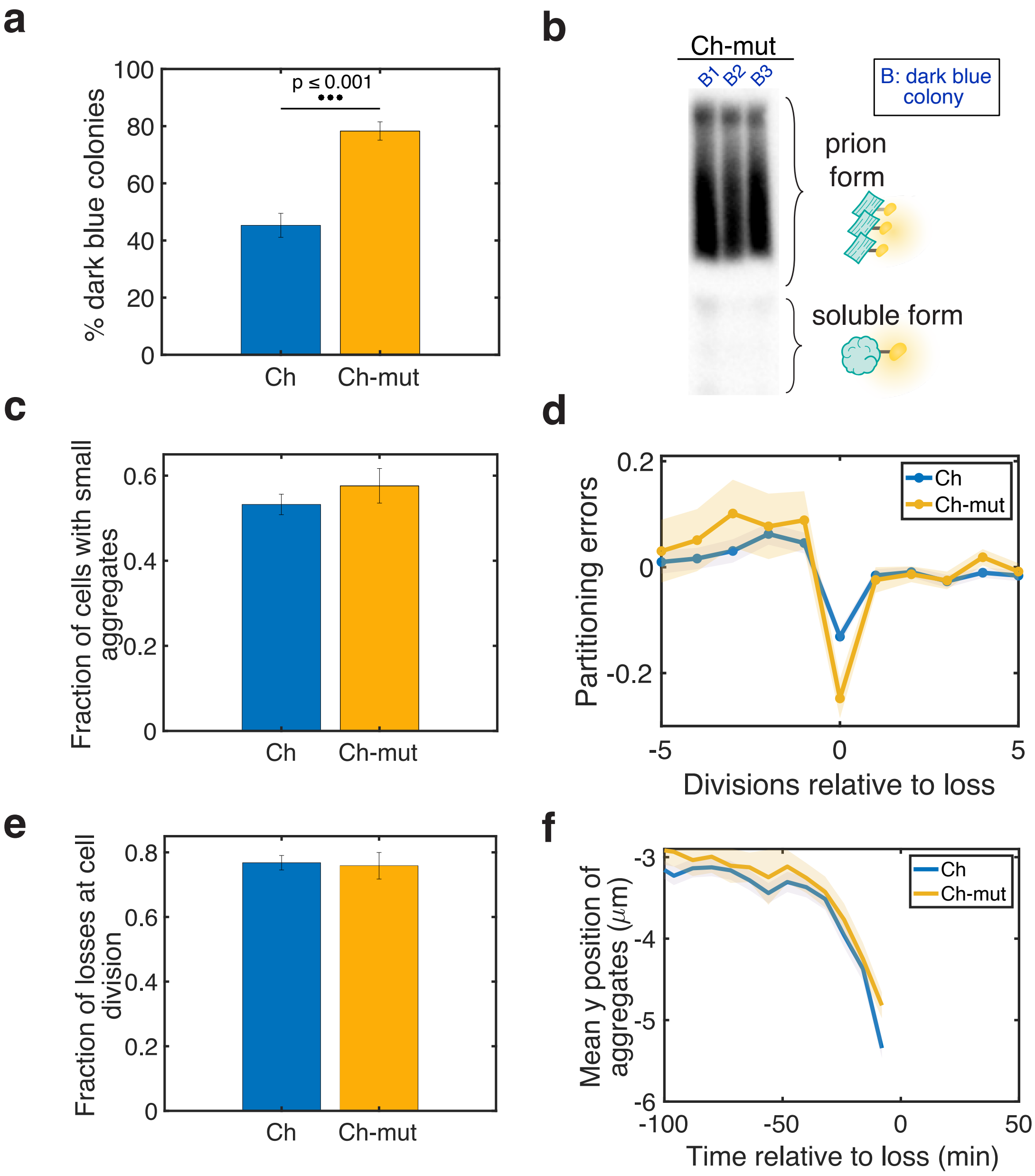

**Fig. S8. A mutant *Ch* SSB PrD (*Ch* SSBmut) forms self-propagating aggregates without exposure to New1 with similar properties to those formed by the *Ch* SSB PrD. a)** Percentage of dark blue colonies (indicating prion-containing cells with the P*clpB-lacZ* assay) upon replating single blue colonies shows that the mutant has a higher fraction of colonies with prion-containing cells (p = 4.1e-04). **b)** SDD-AGE of 3 colonies of the mutant shows the presence of insoluble aggregates. **c-f)** The mutant domain exhibits similar properties to the WT domain as assayed by our microfluidic assay. The fraction of cells with small aggregates (**c**, n = 1,779 cells for the WT and 578 cells for the mutant), the partitioning errors relative to the loss (**e**, n = 349 cells for the WT and 125 cells for the mutant), the fraction of losses at cell division (**e**, n = 754 loss events for the WT and 207 loss events for the mutant), and the position of the aggregates (**f**, n = 754 cells for the WT and 207 cells for the mutant) are similar for both PrDs.

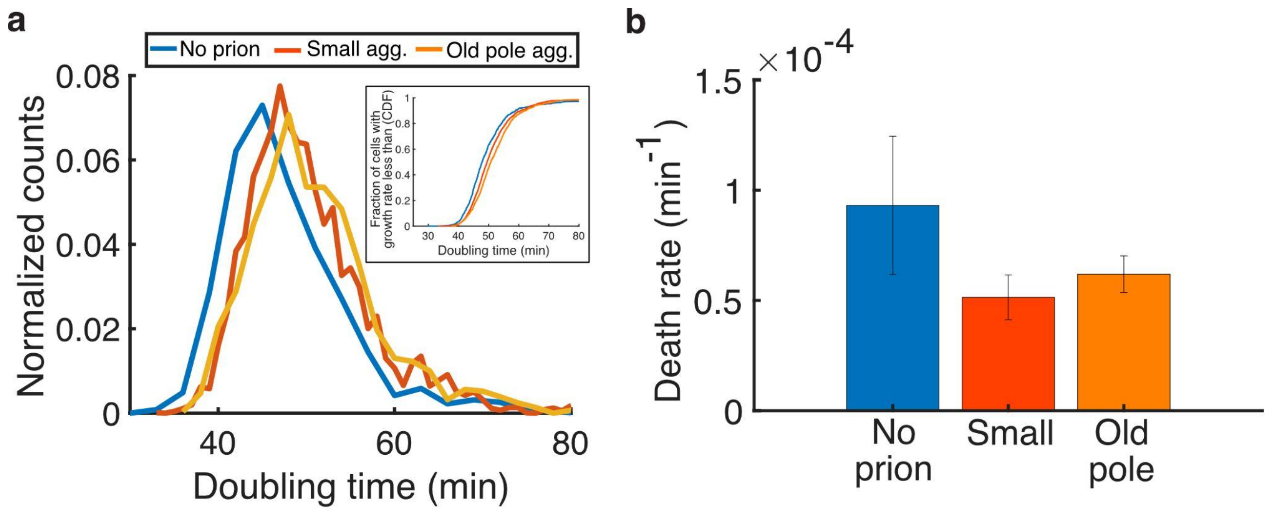

**Fig. S9. Propagation of the *Ch* SSB prion imposes a slight growth rate penalty on the cells. a)** Histogram of estimated doubling time in the microfluidic device for cells without the prion (blue), with the small aggregates (red), and with the old-pole aggregates (yellow, n = 6,767 time points without the prion, 14,154 with small aggregates, and 5,952 with old pole aggregates). The median doubling time is ∼1.5 min greater for cells with small aggregates and ∼3 min greater for cells with old-pole aggregates as compared to cells without prions. (Inset) Cumulative distribution function (CDF) of the doubling time. **b)** Estimated death rate in the microfluidic device of cells without prions (blue), with small aggregates (red), and with old pole aggregates (yellow, n = 8 cells deaths without the prion, 23 with small aggregates, and 50 with old pole aggregates out of 10,740 min, 55,936 min, and 100,943 min observed, respectively)

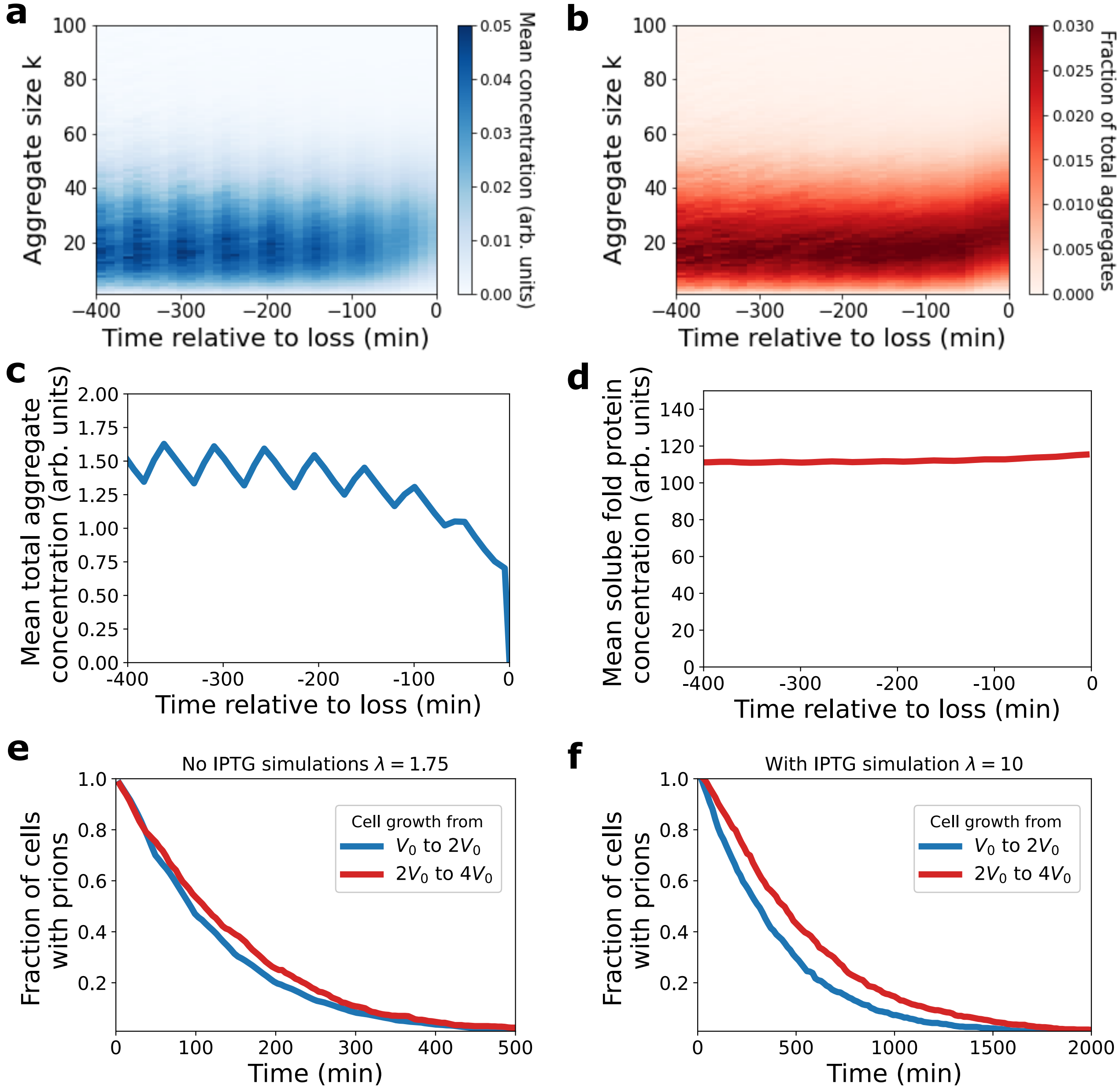

**Fig. S10. The distribution of aggregate size reaches a quasi-stationary state before the time of prion loss in the simulations of our stochastic model. a)** The distribution of aggregate concentrations is cyclo-stationary prior to 200 minutes before loss. The division time here is 50 min. The y-axis corresponds to the aggregate size $k$ where $k$ is the subscript in $Y_k$. Time is binned in bins of 10 minutes, and average concentrations are computed in each bin taken over 100,000 simulations. **b)** The distribution of the fraction of total aggregates is constant prior to loss except for a slight growth in aggregate size right before the loss. The fraction of $Y_k$ was computed in each time bin $i$ as $\langle y_k^i \rangle / \sum_j \langle y_j^i \rangle$. **c)** The mean total aggregate concentration, $\sum_k y_k/V$, is cyclo-stationary prior to 200 min before loss. Aligning the cells at the time of loss effectively aligns the division times because in the model the division time is constant. Mother cells that keep the prion at division tend to gain prion concentration at division because the partition is on average positive, see Fig. 5i of the main text. **d)** The average soluble fold protein concentration $\langle X/V \rangle$ is constant prior to loss except for a slight increase right before the loss. **e)** The model predicts that the prion is more stable in cells with larger sizes. This is because in order for the concentrations of molecules to be identical between small and large cells, larger cells need to have higher numbers of molecules, thus reducing partitioning errors. Each curve corresponds to 1000 simulations with a fixed cell division time of 50 min. Model parameters were estimated as described in Fig. 5 and the SI 3.2 to model the mother machine experiments where there is no IPTG. **f)** In the replating experiments, cells are grown in 10 $\mu$M IPTG. In the model, this corresponds to an increase in the soluble fold protein production parameter $\lambda$. Setting $\lambda = 10$, we find that the difference in prion stability is more stark between small cells and large cells.

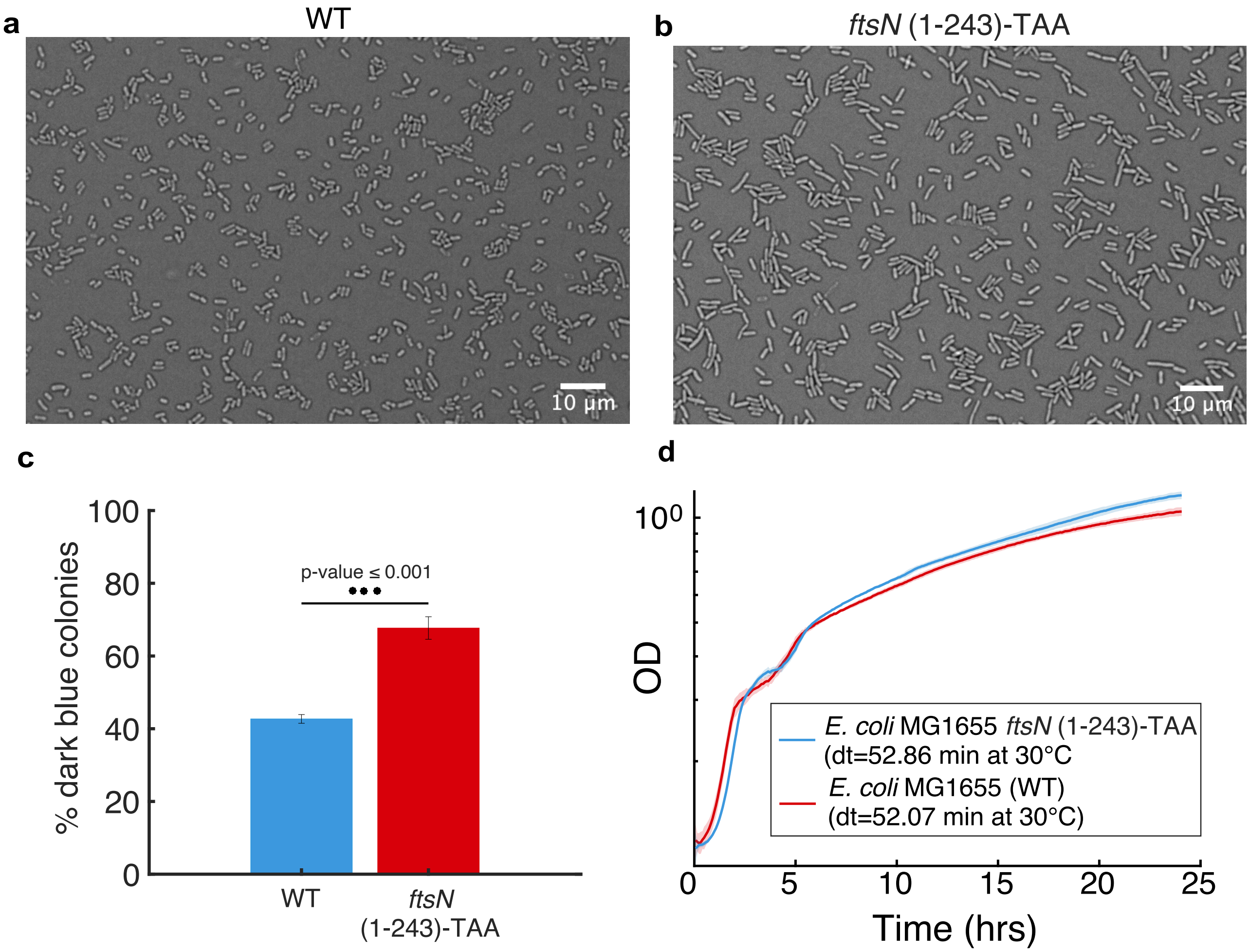

**Fig. S11. A mutant with bigger cell size and similar growth rate propagates the *Ch* SSB prion with a higher stability. a)** 60x microscopy image of background strain BLS80 cells in exponential phase. **b)** 60x microscopy image of BLS333 cells (N-terminal truncation of *ftsN*) in exponential phase, showing elongated cells compared to BLS80 (as previously reported (1)). **c)** The mutant strain with bigger cell size propagates the prion with a higher stability during replating. A pool of colonies containing *Ch* SSB prion aggregates were replated to assay stability and assessed for the presence of the prion using the P*clpB-lacZ* assay (dark blue indicates the presence of the prion). The percentage of colonies with prion-containing cells is higher in the mutant than in the WT, where ∼60% of WT cells lose the prion during the replating compared to ∼35% for the mutant. **d)** Growth curves for the WT (red) and *ftsN* mutant (blue) at 30°C, showing that they grow with similar doubling times (52.1 min for the WT and 52.9 min for the mutant).

### 2. Materials and Methods

#### 2.1. Strains, plasmids, and growth conditions.

***2.1.1. Bacterial strains and growth conditions.*** A complete list of the *Escherichia coli* strains and plasmids can be found in Table S1. The MG1655 strains used for the microfluidic experiments had a motility deletion ($\Delta motA$), expressed a constitutive fluorescent marker for segmentation and tracking (P*rpsL*-mSCFP3A), and contained the P*clpB*-*lacZ* reporter on the F' plasmid for identification of colonies with cells containing prions.

*E. coli* strain NEB® 5-alpha F'IQ (NEB, MA, USA) was used for routine cloning procedures and strains BLS80 and BLS333 were used for prion propagation experiments. Chemically competent *E. coli* were transformed with plasmid DNA by the standard heat shock procedure. *E. coli* strains were grown in LB medium (1% tryptone, 0.5% yeast extract, 1%NaCl) with aeration at 250 rpm or solid media with the appropriate antibiotics at standard concentrations: kanamycin 20 $\mu$g/ml (Km), carbenicillin 100 $\mu$g/ml (Carb), chloramphenicol 12.5 $\mu$g/ml (Cm). All antibiotics and LB medium were sourced from Fisher Scientific.

BLS80 was created in several steps. First, $\Delta lacIYZA$::Km$^R$ was transduced from TB12 (2) (a gift from Thomas Bernhardt; Harvard Medical School, Boston, MA, USA) into LPT37 (3) by P1 phage transduction according to a protocol from Robert Sauer (Massachusetts Institute of Technology; protocol available at:https://openwetware.org/wiki/Sauer:P1vir_phage_transduction), generating strain BLS63. The deletion of *lacIYZA* in BLS63 was verified by colony PCR (using internal primers oBLS107+oBLS108 targeting *lacZ* to test for absence of *lacZ* and primers oBLS109+oBLS110 targeting *motA* as a negative control reaction and primers oBLS138+oBLS139 targeting *cyaA* as a positive control reaction). Absence of *lacIYZA* was additionally confirmed by plating BLS63 on LB plates containing X-gal (40 $\mu$g/ml). Only cells that have lost the *lacIYZA* DNA section would remain pale under these conditions. Next, plasmid pCP20 (encoding the yeast Flp recombinase) was transformed into BLS63 and grown overnight at 30°C on LB Carb plates. The next day, single colonies were streaked on plain LB plates and incubated over night at 42°C. Then, single colonies were picked and re-streaked on LB plates containing either Carb and Km, or no antibiotic and grown overnight at 30°C. A single Carb and Km-sensitive colony was picked and reverified by streaking on the same growth plates and then designated BLS79. Then, pSC101$^{TS}$-empty was transformed into BLS79, grown overnight on LB Cm at 30°C. Finally, the F' containing the $P_{clpB}$-*lacZ* reporter was introduced into BLS79 by mating with strain DH5$\alpha$Z1 F' $P_{\mathrm{clpB}}$-*lacZ* (as described previously(4)), generating BLS80.

BLS333 was also generated in a stepwise manner. Initially, *ftsN* (1-243)-TAA::Km$^R$ was transduced from strain MT190 into BLS79 by P1 transduction as described above, generating strain BLS305. The Km$^R$ gene was flipped out using the pCP20 plasmid as described above, generating strain BLS328. Proper deletion of the 3' codons 244–319 was verified by colony PCR of 12 individual BLS328 clones using primers oBLS387+oBLS388 followed by Sanger sequencing of the amplified DNA products. A single positive BLS328 clone was then used and the F' $P_{\mathrm{clpB}}$-*lacZ* reporter was integrated into this strain by mating as described above, generating strain BLS333.

***2.1.2. Plasmid construction.*** All plasmids generated in this study (see Supplementary Table S1) were constructed using Gibson assembly (NEBuilder® HiFi DNA Assembly Master Mix, NEB, MA, USA) or standard restriction enzyme-based cloning and subsequent ligation using T4 DNA ligase (NEB, MA, USA). Reactions were performed according to the manufacturer's instructions, and Gibson assembly reactions were routinely incubated for 25 min at 50°C. Primers used for plasmid construction are shown in Supplementary Table S2. Sequence integrity was verified by Sanger sequencing by Quintara Biosciences (MA, USA).

pBLS26: *mScarlet-I* was amplified with primers oBLS34+oBLS35 and then ligated into NotI/XbaI-digested pSC101$^{\text{TS-}}$*NEW1* by Gibson assembly.

Plasmids A6 and A9: *E. coli* codon-optimized SSB cPrD DNA sequences for *Lactobacillus heilongjiangensis* and *Moraxella lincolnii* were ordered from Integrated DNA Technologies (IA, USA) containing a 5' NotI sequence followed by a single adenine and 3' TAA stop codon followed by an XbaI site. At the 5' and 3' end, both sequences contained random DNA sequences used for PCR amplification using primers oEF35+oEF432. Following amplification, PCR products were digested by NotI/XbaI (NEB, MA, USA) and then ligated into NotI/XbaI-digested pEF130 using T4 DNA ligase.

pEM04: pEM04 was identified as encoding a *NEW1*-independent *Ch* SSB cPrD mutant during a screen of a *Ch* SSB cPrD library of mutants. This library was created by error-prone PCR using the Diversify PCR Random Mutagenesis Kit (Takara Bio, CA, USA) with primers oEF259 and oEF260, and WT *Ch* SSB cPrD sequence as template. The mutagenized PCR product was ligated into linearized pEF171 via NotI/XbaI sites. The resulting plasmid was transformed into six aliquots of NEB® 5-alpha F'IQ using standard heat shock procedures. After the initial recovery outgrowth, transformants were pooled, plated on selective plates (Carb), and allowed to grow for two days at 30°C. The plasmid was isolated from pooled transformants using standard procedures. To screen for *NEW1*-independent *Ch* SSB cPrD mutants, the library was transformed using standard heat shock procedures into DH5$\alpha$Z1 carrying the $P_{\text{clpB}}$-*lacZ* reporter on the F' but lacking pSC101$^{\text{TS-}}$*NEW1*. Transformants were initially recovered for 2 hours at 30°C, and subsequently diluted into selective media and incubated for a further 23 hours. Finally, transformants were plated on LB plates containing X-gal (40 $\mu$g/ml), Carb, Kn, and IPTG (for induction of *Ch* SSB PrD) and screened for dark blue colonies as described previously (5). Dark blue colonies, indicative of *NEW1*-independent conversion, were selected for plasmid isolation and Sanger sequencing to identify the mutant *Ch* SSB PrD responsible for the dark blue phenotype. One such mutant PrD which we termed SSBmut, encodes the following codon changes: F9Y, N20I, F21Y, N22I and N28F.

Gene block sequences

Lactobacillus heilongjiangensis SSB cPrD (A6 construct):

SGSGNNYSNNNQAPSYNNSNQSNQSPVNNNSNNNFNGGNNAPANSGNYNSNTNNNNNNQSNNNNSSS

Moraxella lincolnii SSB cPrD (A9 construct):

NQNHQGGYQSGYQGGHQGYGNFQNQNGNPNGNQSGFQNHQNTANAYQNNNNNNNFNNNNNNNNFNNTDNHSA
QQAGQNNQNQPKPNN

#### *2.1.3. List of strains and plasmids.*

| Strain | Details | Reference |
|---|---|---|
| NEB® 5-alpha F‘IQ | F‘ *proA$^+$B$^+$ lacI$^q$* $\Delta$*(lacZ)M15 zzf::Tn10* (Tet$^R$) / *fhuA2*$\Delta$*(argF-lacZ)*U169 *phoA glnV44* $\Phi$*80*$\Delta$*(lacZ)M15 gyrA96 recA1 relA1 endA1 thi-1 hsdR17* | New England Biolabs (NEB, USA) |
| DH5$\alpha$Z1 | *lacIQ*, P$_{N25}$-*tetR*, Sp$^R$, *deoR*, *supE44*, $\Delta$(*lacZYA*-argFV169), Phi80 *lacZ*$\Delta$M15, *hsdR17*(rK- mK+), *recA1*, *endA1*, *gyrA96*, *thi-1*, *relA1* | Purchased from Expressys (Rolf Lutz) |
| TB12 | MG1655, $\Delta$*lacIYZA*::Km$^R$ | (2) |
| MT190 | TB28, *ftsN* (1-243)-TAA::Kan$^R$ (*ftsN* deleted of amino acids residues 244–319) | (1) |
| LPT37 | K-12 F– $\lambda$– ilvG– rfb-50 rph-1 $\Delta$*motA*, attTn7::P$_{rpsL}$-*mSCFP3* | (3) |
| BLS63 | LPT37, $\Delta$*lacIYZA*::Km$^R$ | This study |
| BLS79 | BLS63, $\Delta$*lacIYZA*::frt | This study |
| BLS80 | BLS79 with F‘ carrying the P$_{clpB}$-*lacZ* reporter (described in (5)) | This study |
| BLS305 | BLS79, *ftsN* (1-243)-TAA::Km$^R$ | This study |
| BLS328 | BLS305, *ftsN* (1-243)-TAA::frt | This study |
| BLS333 | BLS328 with F‘ carrying the P$_{clpB}$-*lacZ* reporter | This study |

| Plasmids | Details | Reference |
|---|---|---|
| pSC101$^{TS}$-*NEW1* | Temperature-sensitive plasmid expressing NEW1-CFP from P$_{tac}$ | (6) |
| pSC101$^{TS}$-empty | Empty temperature-sensitive plasmid | (6) |
| pCP20 | Temperature-sensitive plasmid containing yeast Flp recombinase | (7) |
| pBLS26 | pSC101$^{TS}$-*NEW1* with NEW1 translationally fused to *mScarlet-I* | This study |
| pEF130 | pBR322-P$_{tac}$-His6-mYFP-*Ch* SSB cPrD | (5) |
| pEF171 | pBR322-P$_{tac}$-His6-mYFP with NotI and XbaI cloning sites at the 5’ end of mYFP to in-frame fuse genes of interest. | This study |
| pEM04 | pBR322-P$_{tac}$-His6-mYFP-*Ch* SSB cPrD carrying F9Y, N20I, F21Y, N22I and N28F mutations in the SSB cPrD sequence. | This study |
| A6 | pBR322-P$_{tac}$-His6-mYFP-*Lh* SSB cPrD | This study |
| A9 | pBR322-P$_{tac}$-His6-mYFP-*Ml* SSB cPrD | This study |

**Table S1. Strains and plasmids. cPrD: candidate prion-forming domain; SSB: Single stranded DNA binding protein; Km$^R$: kanamycin resistance, Tet$^R$: tetracycline resistance, Sp$^R$: spectinomycin resistance.**

#### *2.1.4. List of oligos.*

| Name | Sequence (5'->3') | Purpose |
|---|---|---|
| oBLS34 | GGGCTATCAAGCGGCCGCAGTTTCTAAAGGTGAAGCAGTTATCAAGG | Cloning pBLS26 |
| oBLS35 | GGGGATCTCTCGAGTCTAGATTACTTATACAGTTCATCCATACCTCCGG | |
| oEF259 | ATGAGCTCTACAAAGCGGCCGCA | Construction of mutant *Ch* SSB PrD library, including pEM04 |
| oEF260 | AGAGGCCCCAAGGGGTTATGCTATCTAGATTA | |
| oEF35 | CATCGCCGCTTCCACTTT | Cloning A6 and A9 |
| oEF432 | GTGTCGCCCTTATTCCCT | |
| oBLS107 | GTTTTACAACGTCGTGACTGGG | Verification of $\Delta$*lacIYZA*::Km$^R$ |
| oBLS108 | GATAACTGCCGTCACTCCAGC | |
| oBLS109 | GATTAAAGGCACGCTGAAGGC | |
| oBLS110 | AAGCAGAGTGACTTTGACGC | |
| oBLS138 | CGCCTGATGAAACTCAACGCC | |
| oBLS139 | CGCGTTCACGGCTGAGCTTTTC | |
| oBLS387 | AGCGCCAACGTCAGGCGC | Verification of *ftsN* (1-243)-TAA |
| oBLS388 | CGATGACCACATGGCCGTTACG | |

**Table S2. Oligonucleotides.**

**2.2. Prion induction and LacZ assay.** The strain BLS80 was made chemically competent following the transformation storage solution (TSS) method. To induce prion formation, the pBLS26 (New1) and pEF130 (*Ch* SSB mYFP) plasmids were co-transformed into the TSS competent cells (heatshock at 42°C for 30 seconds, incubate on ice for 2 minutes) with an extended recovery time of two hours at 30°C before plating on LB agar plates containing the appropriate antibiotics. After overnight growth at 30°C, a colony selected from the transformation plate was grown in liquid LB containing antibiotics and 10$\mu$M IPTG (Fisher Scientific) at 30°C. By simultaneously producing New1 and the cPrD fusion protein, prion formation was induced. After 16 hours, the cells were plated on indicator plates: LB agar containing 10 $\mu$M IPTG, X-gal (40 $\mu$g/mL), Phenylethyl-$\beta$-D-thiogalactopyranoside (TPEG) (Gold Biotechnology) (500 $\mu$M), Km and Carb at 37°C for 24 hours to cure the New1 plasmid (temperature sensitive). As all cells express ClpB to some extent, TPEG was included in the plates as a competitive inhibitor of the enzyme $\beta$-galactosidase, making it easier to distinguish the colonies with prion-containing cells (dark blue, which express ClpB at a higher level) from the colonies without prion-containing cells (pale blue).

**2.3. Verification of curing.** To verify the curing of New1 after incubation at 37°C, the prion-containing cultures were inspected under the microscope after 16h of growth. An agar pad with 1 $\mu$l of culture was used to visualize the cells. Blue (constitutive CFP), yellow (SSB mYFP plasmid) and red (New1-mScarlet plasmid) fluorescence channels were used (**Fig. S1a**). When no red fluorescence was detected (**Fig. S1b**), it was determined that curing of New1 was successful. To corroborate these findings, prion-containing cultures were plated in LB plates with and without chloramphenicol. These plates were incubated at 30°C for 24h. The curing was determined to be complete since no colonies were observed with chloramphenicol (**Fig. S1c**).

**2.4. Re-plating experiments.** To confirm prion propagation, single prion-containing colonies (dark blue) and prion-free colonies (pale blue) were re-suspended, serially diluted, re-plated on indicator medium, and incubated at 30°C for 30h. This process was repeated generating Round 1 (R1), R2, and R3 plates (**Figs. 3c, S8a, S11c**). At each round, all colonies were counted (dark and pale blue) and the percentage of dark blue colonies was determined. To determine propagation dynamics independent of individual colonies (lineages), we performed pooling experiments (**Fig. S7a**). Here, a mixture of dark blue colonies was re-suspended. The subsequent rounds and calculations were generated as mentioned above.

For the *Ch* SSB PrD stable lineage, microfluidic experiments were performed at each round of re-plating (**Fig. 4b**) to observe propagation dynamics over time. In this case, individual dark blue colonies were cultured and induced overnight in liquid media with the appropriate antibiotics and 10 $\mu$M IPTG. These cultures were serially diluted, re-plated on indicator medium, and incubated at 30°$C$ for 30h, giving rise to R1 plates. Subsequent rounds were generated repeating this process.

### 2.5. Microfluidic experiments and microscopy.

#### *2.5.1. Microfluidic device.*

**2.5.1.1 Fabrication of the wafers.** The molds for the microfluidic device used in this study were fabricated at the McGill Nanotools Microfabrication Facility (Montréal, Canada) using standard photolithography practices. The molds consist of two layers built up on a silicon wafer, with the first layer containing the cell trenches and the second layer making up the main feeding channel. A 4-inch silicon wafer was washed with acetone, isopropyl alcohol (IPA) and deionized water, and then dehydrated at 150°C for 15 minutes. The wafer was then cleaned with oxygen plasma in the DSB6000 Oxygen Asher. Photoresist (5 mL SU-8 2001, Microchem) was dispersed across the wafer at 500/87/10 (rpm/acceleration/time), and then at 1500/348/60 (rpm/acceleration/time) using a Laurell Spin Coater. The wafer was then soft baked (SB) for 1 minute at 65°C, 3 minutes at 95°C, 1 minute at 65°C before exposure to UV light (42.5 $mW/cm^2$) for 1.5 seconds using the EVG620 photomask aligner. UV exposure was followed by a post exposure bake (PEB) of 1 minute at 65°C, 20 minutes at 95°C, 1 minute at 65°C. Excess SU-8 was washed away by swirling gently in a dish filled with SU-8 developer for 30 seconds, then rinsed immediately with more developer, followed by IPA for 10 seconds. After the final hard bake (HB) for 1 minute at 65°C, 10 minutes at 150°C, 1 minute at 65°C, the height of the first layer was measured using the Ambios XP200 profiler. A second coating of photoresist (5mL SU-8 2015, Microchem) was applied, spun first at 500/87/10 (rpm/acceleration/time), and then 3000/348/60 (rpm/acceleration/time) using the Laurell Spin Coater. SB for 1 minute at 65°C, 4 minutes at 95°C, 1 minute at 65°C. A developer soaked swab was used to clean off the photoresist from the alignment markers, followed by a second SB for 1 minute at 65°C, 4 minutes at 95°C, 1 minute at 65°. The second photomask was aligned with the features from the first layer on the wafer and exposed to UV light (42.5 $mW/cm^2$) for 3.4 seconds (EVG620 photomask aligner). PEB was done for 1 minute at 65°C, 4 minutes at 95°C, 1 minute at 65°C. Excess photoresist was washed away by swirling gently in a dish filled with SU-8 developer for 1 minute and 30 seconds, then rinsed immediately with more developer followed by addition of IPA for 10 seconds. A final HB step was done for 1 minute at 65°C, 15 minutes at 150°C, 1 minute at 65°C.

**2.5.1.2 Preparation of the device.** Polydimethylsiloxane (PDMS) (Sylgard 184 Silicon Elastomer, Fisher Scientific) was mixed at a 10:1 (monomer:curing agent) ratio, poured on top of a 1.0 $\mu$m tall wafer and degassed for one hour at room temperature before baking for an additional 1.5 hours at 65°C. After careful removal of the PDMS from the wafer, individual PDMS chips were cut out with a razor blade. The inlet and outlet holes were punched with a 0.75 mm biopsy puncher (World Precision Instruments). The PDMS chips were sonicated in isopropyl alcohol (Fisher Scientific) for 30 minutes and dried at 65°C for 15 minutes. Glass coverslips (Fisher Scientific: 22x40 mm #1.5) were cleaned with 1M potassium hydroxide (KOH, Sigma Aldrich) for 20 minutes. The PDMS chips were bonded to the glass coverslips using a plasma cleaner (Oxygen flow rate at 45 sccm, power at 30W for 15 seconds, Tergeo Plasma Cleaner, PIE Scientific). The completed microfluidic device was heated to reinforce the plasma bonding at 100°C for 10 minutes, then 65°C for 30 minutes.

***2.5.2. Microscopy experiment.*** Typically around 20 prion-containing colonies (dark blue from the *lacZ* assay) were individually cultured for 16 hours at 30°C in 5mL of imaging medium (M9 salts (Sigma Aldrich), 0.2% (w/v) glucose (Alfa Aestar), 1 mM $MgSO_4$ (Sigma Aldrich), 0.1 mM $CaCl_2$ (Sigma Aldrich), 20$\mu$g/mL uracil (Sigma Aldrich), 0.2 g/L casamino acids (Bacto Casaminoacids, Thermo Fisher Scientific) and 0.85 g/L Pluronic F-108 (Sigma Aldrich) supplemented with Km, Carb, and 10 $\mu$M IPTG. Each culture was inspected using fluorescent microscopy to confirm the presence of prions and the absence of New1. After prion content was confirmed, cells were loaded into the main feeding channel of a microfluidic chip and then centrifuged into the cell trenches (5000g for 10 minutes). A syringe pump (New Era Pump System) was connected to the inlet hole via flexible plastic tubing (Tygon) to provide the cells with fresh imaging medium (without antibiotic and without IPTG) at 5 $\mu$l/min for the duration of the experiment.

All microscopy was conducted using a Zeiss Axio Observer inverted microscope equipped with a 63x Plan-Apochromat M27 oil objective (NA 1.40), an Orca Flash 4.0 LT camera (Hamamatsu), and an LED epifluorescence illuminator (Colibri 7), inside a temperature controlled incubation chamber set at 30°C. Exposure time (100 ms) and light intensity (10-20%) were low to reduce photobleaching, with 16-bit CZI images taken every 8 minutes using the Colibri 7 (Zeiss) 91 HE filter set. Focal drift was corrected automatically with the Definite Focus 2 (Zeiss) monitoring and compensation system using an infrared laser (850 nm).

#### 2.5.3. Data processing.

**2.5.3.1 Segmentation and lineage tracking.** Image segmentation and single-cell trace assembly were done according to a previously described method (8), in which a pixel mask is created to define the boundaries of each cell based on the constitutively expressed CFP. Within the pixel mask, data (cell size, fluorescence, etc.) were extracted and the mother cells from consecutive frames were concatenated based on matching centroid position, creating a continuous time trace for each individual cell. We define average fluorescence as the mean pixel value (YFP or CFP) within the cell's segmentation pixel mask, and peak fluorescence as the median value of the brightest 10% of pixels within the segmentation pixel mask. We identified cell division by a decrease of cell area to less than 60% of its previous value. The traces were then manually curated to remove dead cells and obvious segmentation errors.

**2.5.3.2 Equilibration of growth conditions.** To maximize the number of prion-containing cells in the microfluidic experiments, the cells were grown in 10 $\mu$M IPTG until they were loaded in the microfluidic experiments (see Sec. 2.2 and 2.5.2). To keep the protein levels as low as possible, IPTG was not supplemented in the imaging media for the duration of the experiment, relying instead on leaky expression of the promoter. Therefore, the YFP fluorescence decreased at the beginning of the experiment to adjust to the new growth conditions (**Fig. S2a**). We thus started the analysis of the data after the YFP reached the equilibrium, as shown in **Fig. S2a-c**.

**2.5.3.3 Prion loss and spot finding.**

**2.5.3.3.1 Spot tracking** Spots were identified within cells by finding peaks of YFP fluorescence using the local maxima, which produced an (x, y) value for the center of a spot along with the number of spots present in a cell. We used the peak finding open source function 'findpeaks' (Nathaniel C. Yoder), that uses a low pass Gaussian filter (5x5 pixel size, standard deviation of 0.95) to emphasize point spread function-size features. Cells were defined as starting with the prion if aggregates were detected in at least 3 of the first 15 time points of the experiment. A prion loss event was defined as the first time point in a run of at least 8 consecutive images without any spots. If a spot was detected after the loss, the entire trace was excluded from the analysis. These rare traces were typically found to contain segmentation errors and pixels from other cells.

**2.5.3.3.2 Prion loss curve** The prion loss curve (PLC), or the fraction of tracked cells with prions, was estimated according to the definition of the cumulative distribution function (CDF) of prion propagation duration $T$.

$$\begin{aligned}\mathrm{PLC}(t) &= 1 - \mathrm{CDF} = 1 - P(T \leq t) = P(T > t) \\ &= \frac{\text{number of cells that propagated the prion for longer than } t}{\text{number of cells that started with the prion that were tracked longer than } t}.\end{aligned}$$

We used this definition as cells were sometimes lost in the tracking or washed out of the device before they lost the prion, and their exact duration of propagation was unknown. However, they could still be included in the calculation since they propagated the prion until a certain time point $t_0$, and their duration of propagation is therefore greater than $t_0$.

**2.5.3.3.3 Aggregate classification** Two different methods were used to classify the types of aggregates which gave similar results (**Fig. S4a-b**). The first method is based on the mobility of the aggregates, where the goal was to identify aggregates that were largely immobile. We calculate the minimum possible displacement between time points for each aggregate. If more than one aggregate was detected in particular time points, we calculated the minimum possible movement for each aggregate, and then took the smallest movement at each time point. We then calculated the moving average over periods of $\sim$3 cell division (18 time points of 8 min) for each time trace, and calculated the minimum for each cell (**Fig. S4a**). This showed a good separation between largely immobile aggregates with average movement of $\sim$ 1 pixel, and mobile aggregates with an average movement of $\sim$ 5 pixels. Cells with a minimum displacement below 2.5 pixels were classified as old-pole aggregates-containing cells, and cells with larger displacements as small aggregates-containing cells.

The second method relied on the brightness of the old-pole aggregates. A ratio of the median top 10% of YFP pixel values in the cell image (peak) divided by the YFP protein concentration (average) for the entire cell was used to classify the types of aggregates (**Fig. S4a**). The median ratio was calculated for each trace, and a value of above 5 (peak/average) was classified as old-pole aggregates-containing cells whereas below this value the cells were classified as small aggregates.

These two methods gave overall similar results (**Fig. S4b**). The minor difference was explained by a small number of cells that had aggregates that were mostly immobile and localized to the old pole yet were not very bright. These were classified as old-pole using the mobility methods, but small aggregates using the brightness methods. These cells were very stable, and showed long-term stability in the "small aggregates" using the brightness method (**Fig. S4b**). Therefore, we used the mobility method throughout the paper, since it classifies the cells according to the phenomenological description of immobile aggregates.

**2.5.3.4 Fraction of losses in one of the two daughter cells.** To estimate how frequently both the mother and daughter cell lost the prion simultaneously, we selected the loss events in the mother that contained tracking information for the cell right below (the daughter cell). Since our tracking software only contains information for the top two cells in the trenches, we used time points for the second cell until the mother divided after the loss (i.e. the "daughter cell"), up to a maximum of 6 time points. Daughter cells that had at least 3 time points were kept for analysis, and if at least two time points contained detected aggregates these cells were considered to have kept the prion.

**2.5.3.5 Partitioning errors.** To calculate the partitioning errors, we selected the cell divisions that contained tracking information for both the mother (cell tracked for the duration of the experiment) and daughter cell (cell immediately below). We curated cell divisions where the difference in cell area between the newborn was less than 25% of the mother cell to avoid possible segmentation errors, and selected cells whose fluorescence was less than 10 times the population mean to avoid outliers. The partitioning errors were then calculated between the two newborn cells according to this formula:

$$\text{Partitioning error} = \frac{\text{YFP}_{\text{mother}} - \text{YFP}_{\text{daughter}}}{\text{YFP}_{\text{mother}} + \text{YFP}_{\text{daughter}}}$$

**2.5.3.6 Doubling time and death rates.** The doubling time was estimated at each cell division according to the following formula:

$$\tau_{\text{doubling}} = \frac{t_f - t_i}{\log_2 \left( A(t_f)/A(t_i) \right)}$$

Where $A(t)$ is the cell area, $t_i$ the first time point of the cell cycle, and $t_f$ is the last point of the cell cycle before cell division. The death rate was calculated using the uncurated data that includes cells that stopped growing (which would have been excluded from the other analyses). Cells were considered dead if they were growing at the beginning of the experiment, and their average doubling rate ($g_2 = 1/\tau_{\text{doubling}}$) during the last 5 time points of the experiment was less than 20% of the population average. Their time of death was estimated as the first time their growth rate was below 20% of the population average. Cells whose growth slowed below that threshold and increased again were rejected from further analysis.

**2.5.3.7 Bootstrapping.** Where indicated, the standard error on the measurement was estimated with bootstrapping. For an experiment containing $n$ lineages/time traces, $n$ lineages were picked randomly with replacement from the original set. The measurement was calculated using this randomly picked set of cells, and this procedure was repeated 100 times. The standard deviation was calculated across these 100 repeats.

**2.6. SDD-AGE.** For SDD-AGE analysis, plasmids pEF130, A6, A9 and pEM04 were co-transformed into BLS80 together with pBLS26 and plated on LB plates containing Carb, Km, Cm and grown overnight at 30 °C. The next day, individual colonies were picked and grown as "Starter Cultures" (SC) at 30 °C in 800 $\mu$l of LB Carb, Km, Cm and 10 $\mu$M IPTG (IPTG). After about 18 h growth, OD600 values were recorded and cells were serial diluted to 1:100.000 and plated on indicator plates (LB, Carb, Km, IPTG, X-gal (40 $\mu$g/ml) and then grown overnight at 37 °C to lose the temperature-sensitive NEW1-bearing plasmid. Additionally, 500 $\mu$l cells of the SC were pelleted by centrifugation (1 min, 21,000 × g at room temperature (RT)) and stored at -80 °C until further usage. The next day, dark blue or pale colony counts were recorded and indicated colonies were picked and grown as "Colony Cultures" (CC) in 800 $\mu$l LB Carb, Km, IPTG over night at 30 °C for about 18 hours. Finally, OD600 values of CC were recorded and 500 $\mu$l cells were pelleted by centrifugation (1 min, 21,000 x g at RT) and stored at -80 °C until further usage. Cell lysis of SC and CC cell pellets was essentially performed as described before (5). For this, cell pellets were resuspended in lysis buffer: BugBuster protein extraction reagent (MilliporeSigma, MA, USA) supplemented with 1× cOmplete, EDTA-free protease inhibitor cocktail (MilliporeSigma, MA, USA), 1 U/$\mu$l (final concentration) rLysozyme (MilliporeSigma, MA, USA), and 0.5 U/$\mu$l (final concentration) Benzonase nuclease (MilliporeSigma, MA, USA). The lysis buffer volume was calculated as follows: OD600 × volume (ml) of culture pelleted × 20. Cells were lysed for 30 min at RT and mixed 3:4 in 4x SDD-AGE sample buffer (80 mM Tris-HCl, 40 mM acetic acid, 2 mM EDTA, 20% glycerol, 8% SDS and bromophenol blue). 17 $\mu$l of this mixture were then loaded on a 1.5% TAE-based agarose gel prepared containing 0.1% SDS and run at 160 V for 2.5 hours at 4 °C in TAE containing 0.1% SDS running buffer. After separation, proteins were transferred to a nitrocellulose membrane (Protran NC 0.45 $\mu$m; GE Healthcare Bio-Sciences, PA, USA) using the TURBOBLOTTER$^{\text{TM}}$ downward capillary transfer system (GE Healthcare Bio-Sciences).

**2.7. Western blotting.** Upon over night transfer of the proteins to the nitrocellulose membrane, free binding sites were blocked in blocking buffer (TBS: 50 mM Tris-HCl, 150 mM NaCl, pH 7.4, supplemented with 5% non-fat dry milk) for 30 min at RT and then incubated with mouse anti-His primary antibody (THE™ His Tag Antibody, 1:4000 diluted in blocking buffer; Genscript, NJ, USA) for 1 hour at RT. Blots were washed 4x with TBS and subsequently incubated with horse anti-mouse IgG (HRP-linked) secondary antibody (1:10,000 diluted in blocking buffer; Cell Signaling, MA, USA) for 1 hour at RT. Afterwards, blots were washed 4x with TBS and proteins were visualized using Pico ECL Reagent (ProSignal, Genesee Scientific, CA, USA) in a ChemiDoc MP system (BioRad, CA, USA).

**2.8. Growth curve.** The *E. coli* strains BLS80 and BL333 were grown at 30 °C with aeration in 96 well plates with 200 $\mu$l of LB media with the appropriate antibiotics. A Biotek Synergy H1 microplate reader (Agilent Technologies) was used to determine OD600 measurements every 5 min for 24h. Cultures were done in triplicates and sterile media was used as a blank. Growth curves were graphed using MATLAB. The doubling time during the exponential phase was determined using the following formula:

$$\tau_{\text{doubling}} = \frac{t_2 - t_1}{\log_2\left(\text{OD}(t_2)/\text{OD}(t_1)\right)}$$

### 3. Supplementary Results and Discussion

**3.1. Prion content in colony.** The fraction of cells containing prion aggregates varied from colony to colony, presumably due to the stochastic propagation and loss of the prion during the growth of each colony. To assay this phenomenon, we selected dark blue colonies containing low, medium, and high fraction of cells with aggregated fluorescence (**Fig. S6d**). These colonies were assayed for SDS-insoluble aggregates using SDD-AGE, which showed that colonies with a higher fraction of cells containing aggregated fluorescence had a higher fraction of PrD in the insoluble form (**Fig. 3b**). We thus concluded that the variability observed between dark blue colonies in the SDD-AGE in the fraction of proteins in the insoluble state was due to variability in the fraction of cells containing aggregates in these colonies.

#### 3.2. Stochastic model.

***3.2.1. Definition of the stochastic model.*** In our experiments we follow single mother cells that grow and divide for many divisions. We thus simulated growing and dividing cells, keeping track of a single cell in the lineage. Specifically, we model cellular growth as a cell volume $V$ that grows exponentially between division times, see Fig. S12. At each division time the cell volume is reduced by a factor of 2, and each molecule is either kept or removed with a 50 percent probability. We let the division times occur with constant time intervals and let the growth rate be constant throughout the simulation.

This model of cell growth and division depends on a single parameter, namely the division time (which also sets the growth rate). For the experiment whose data is shown in Fig. 1d-e and Fig. 2 of the main text, we measured an average division time of 49.4 minutes. We thus set the division time in our model to be 50 minutes.

We model the rest of the prion propagating system as a continuous-time Markov process. We denote the soluble fold protein as $X$, with total numbers in the cell given by $x$. Production of soluble fold protein numbers is modelled by a stochastic reaction

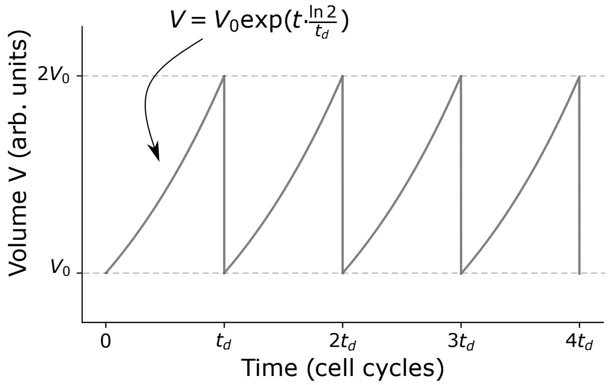

**Fig. S12. A simple model of cellular growth.** We model cellular growth and division as an exponentially growing cell volume that divides at equal time intervals of $t_d$ with symmetric divisions.

with a rate proportional to the cell volume,

$$x \xrightarrow{\lambda V} x + 1. \tag{1}$$

This phenomenological model of protein production makes it so that the $X$ concentration, given by $x/V$, remains on average constant throughout the cell cycle. Specifically, because the numbers grow exponentially between divisions, along with the cell volume, the ratio of numbers to volume remains on average constant. We use this phenomenological model to reproduce the homeostasis of protein concentrations in exponentially growing cells (9).

We denote the prion fold aggregate made up of $k$ proteins as $Y_k$, with total numbers in the cell given by $y_k$. We assume that the $X$ molecules and $Y_k$ aggregates are diffusing around the cell volume and are equally likely to be at any part of the total volume at any given time. We model conversion and elongation as a reaction that occurs when an $X$ molecule and a $Y_k$ aggregate collide. In that case, assuming that the molecules have individual velocities that follow a Boltzman distribution, it can be shown (10) from first principles that the conversion/elongation reaction rate is given by

$$(x,\ y_k) \xrightarrow{\gamma[X][Y_k]V} (x-1,\ y_k+1) \quad \text{for} \quad k = 1, 2, 3 \ldots \tag{2}$$

where $[X] = x/V$ and $[Y_k] = y_k/V$ are the protein and aggregate concentrations respectively. In other words, the protein concentrations follow mass action kinetics, and since $x = [X]\,V$, the rate of production of the total numbers gets scaled by a factor of $V$.

Similarly, we model fragmentation of the prion fold aggregates as a reaction that occurs when a chaperone protein collides with an aggregate. When a collision occurs, we let each binding between any two prion fold proteins to have equal probability of breaking. In that case, the fragmentation reaction is given by

$$(c,\ y_k,\ y_j,\ y_{k-j}) \xrightarrow{\tilde{\alpha}[C][Y_k]V} (c,\ y_k-1,\ y_j+1,\ y_{k-j}+1) \quad \text{for} \quad j = 1, \ldots, k-1, \tag{3}$$

where $[C]$ denotes the concentration of chaperones in the cell. Assuming the chaperone concentration is constant, we define $\alpha := \tilde{\alpha}\,[C]$ resulting in the reaction rate shown in Fig. 5a of the main text.

We assume that random partitioning of molecules at cell division follows a binomial distribution with each molecule having probability 1/2 to be in either daughter cells. In which case, at each cell division time, the following "reaction" occurs

$$x \xrightarrow{\text{At division}} \text{Bin}(x, 1/2) \quad \text{and} \quad y_k \xrightarrow{\text{At division}} \text{Bin}(y_k, 1/2) \quad \text{for} \quad k = 1, 2, 3 \ldots \tag{4}$$

Lastly, we include a minimum seed size in the model in which any aggregate of such size or less is immediately fragmented back into normal fold proteins. Denoting the minimum seed size by $n$, this means that if $k \leq n+1$ and $y_k > 0$ the following immediately occurs

$$(x,\ y_k) \longrightarrow (x + k y_k,\ 0). \tag{5}$$

##### *3.2.2. Simulation procedure.*

**3.2.2.1 The algorithm.** Exact time trajectories of the abundances in the above model can be generated using the Gillespie Algorithm (11), with an additional step to account for cell growth and division (12). The algorithm goes as follows:

**1.** Given the state of the system $(x,\ \{y_k\}_k)$ at time $t_0$, pick the waiting time for the next reaction event to occur. This is done by picking a random number from an exponential distribution with cumulative distribution function $F(t) = 1 - e^{-\int_{t_0}^{t_0+t} r_T(t')dt'}$, where $r_T$ is the total rate of reaction events at time $t$:

$$r_T(t) = \lambda V(t) + \sum_{k=1}^{\infty} \gamma \left(\frac{x}{V(t)}\right)\left(\frac{y_k}{V(t)} V(t)\right) + \sum_{k=1}^{\infty}\sum_{j=1}^{k-1} \alpha \left(\frac{y_j}{V(t)} V(t)\right).$$

However, given the time-dependence of the volume $V(t)$, finding the waiting time this way requires numerically integrating $r_T(t)$ which is computationally time consuming and introduces numerical error. Instead, we can introduce a virtual variable $Z$ that follows the following null rate

$$z \xrightarrow{R_z(t)} z, \tag{6}$$

where $R_z(t) = \max\{r_T(t)\}_V - r_T(t)$ with the rate $r_T$ maximized over all possible volumes

$$\max\{r_T(t)\}_V = \lambda 2 V_0 + \sum_{k=1}^{\infty} \gamma \left(\frac{x}{V_0}\right)\left(\frac{y_k}{V_0} V_0\right) + \sum_{k=1}^{\infty}\sum_{j=1}^{k-1} \alpha \left(\frac{y_j}{V_0} V_0\right).$$

Introducing this variable into the model does not affect the dynamics of $(x,\ \{y_k\}_k)$, and it allows us to use the Gillespie algorithm without having to numerically integrate $r_T(t)$. That is, now the system has a new total reaction rate given by $\tilde{r}_T(t) = r_T(t) + \max\{r_T(t)\}_V - r_T(t) = \max\{r_T(t)\}_V$. This total rate does not have any time-dependence through the volume, and since the abundances $(x,\ \{y_k\}_k)$ do not change by definition of the waiting time in $(t_0, t_0+t)$, the waiting time is picked from the following cumulative distribution function

$$F(t) = 1 - e^{-\int_{t_0}^{t_0+t} \tilde{r}_T(t')(t')dt'} = 1 - e^{-\tilde{r}_T(t_0)t}, \tag{7}$$

where $\tilde{r}_T(t_0) = \max\{r_T(t_0)\}_V$. The waiting time for the next event to occur is obtained by taking a random number from the cumulative distribution function given by Eq. (7).

**2.** Update the cell volume according to the waiting time t:

$$V(t_0+t) = V(t_0) + \int_{t_0}^{t_0+t} V'(t')dt'.$$

**3.** If $V(t_0+t) \geq 2V_0$, then a division has occurred during the waiting time. We first bring the time back to the exact moment when $V$ reaches the threshold $2V_0$: $t_0 \to t_{div}$ where $t_{div}$ is such that $V(t_{div}) = 2V_0$. We then update the cell volume that divides: $V(t_0) \to V(t_0)/2$. We next update the abundances that are reduced due to cell division and random partitioning

$$x \longrightarrow \text{Bin}(x, 1/2) \quad \text{and} \quad y_k \longrightarrow \text{Bin}(y_k, 1/2) \quad \text{for } k = 1, 2, 3\ldots$$

We then go back to step 1.

**4.** If on the other hand $V(t_0+t) < 2V_0$, pick which of the reactions occurs at the time obtained in 1, where the i-th reaction occurs with probability $\frac{r_i}{\tilde{r}_T}$. For instance, an $X$ molecule is converted to prion form and elongates an aggregate of size $j$ with probability $\gamma\,[X]\,[Y_j]\,V/\tilde{r}_T$.

**5.** Update the system according to the reaction that was picked in 4. For instance, if the event turns out to be that an $X$ molecule is converted to prion form and elongates an aggregate of size $j$, then update the system as

$$(x,\ y_j) \longrightarrow (x-1,\ y_j+1).$$

Note that if the null reaction in Eq. (6) is picked, nothing occurs.

**6.** Update the time: $t_0 \to t_0 + t$. We then go back to 1 and re-iterate.

**3.2.2.2 Initial conditions.** An initial condition needs to be chosen to start the Gillespie algorithm. The initial cell volume $V(0)$ in each simulation is chosen to be a random number taken from a uniform distribution that ranges from $V_0$ to $2V_0$. Without loss of generality, we choose volume units such that $V_0 = 1$.

In our experiments cells are grown in 10 $\mu$M IPTG to exponential phase prior to loading into the mother machine. We find that prior to loading, cells with prions typically only had one large aggregate. This might be due to the fact that at exponential phase the chaperones cannot fragment the aggregates as fragmentation is ATP dependent, whereas elongation can still occur as it does not depend on ATP. Once the mother machine experiment begins, the average fluorescence is initially large and eventually reaches an equilibrium, see Fig. S2 and Sec. 2.5.3.2. In the model, the only production of new proteins is given by Eq. (1) with removal of proteins solely driven by cell division. Therefore, the protein concentration will eventually reach a stationary average of $\lambda t_d/\ln(2)$. In Fig. S2 we find that the initial fluorescence is approximately 6 times larger than the equilibrium. We thus set the initial condition to be $x = 0$ and $y_m = 1$ where $m$ is the closest integer to $6V(0)\lambda t_d/\ln(2)$, corresponding to a large initial aggregate that leads to the cell having an initial protein concentration that is 6 times larger than the stationary average. All other $y_k = 0$ when $k \neq m$.

**3.2.2.3 Waiting for equilibrium.** In the experiments, we start our analysis once the concentration of the *Ch* SSB PrD reaches equilibrium, see Fig. S2. This procedure is reproduced in our simulations. In particular, for a given set of model parameters, we run 1000 simulations and compute the average protein concentration, given by

$$[P]_{\text{avg}}(t) = \frac{1}{1000}\sum_{i=1}^{1000}\left(\frac{x^i(t)}{V^i(t)} + \sum_{k=1}^{\infty} k\frac{y_k^i(t)}{V^i(t)}\right), \qquad [8]$$

where $x^i(t)$, $y_k^i$, and $V^i$ correspond to the $x$, $y_k$ numbers and the cell volume from the i-th simulation. To keep track of time numerically we discretize time by taking averages every 10 minutes in simulation time. We then start the analysis once the average protein concentration is within 2.5% of the stationary average given by $\lambda t_d/\ln(2)$, see Fig. S13.

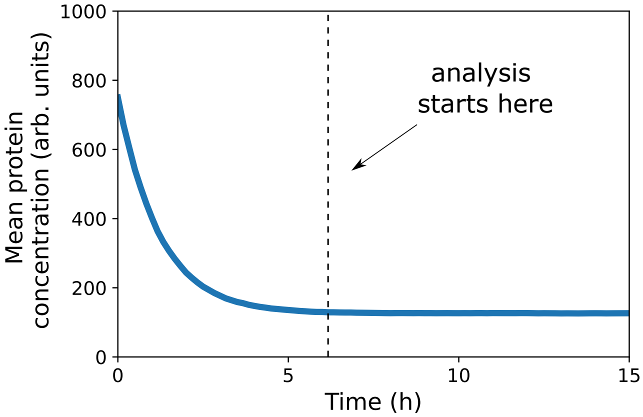

**Fig. S13. Equilibrium of the mean protein concentration prior to beginning of the analysis.** Plotted is the mean protein concentration given by Eq. (8) taken over 1000 simulations. Each simulated cell starts with a large initial aggregate as described in Sec. 3.2.2.2. We start the analysis once time has reached the waiting time, defined as the time it takes for the mean protein concentration to decay to within 2.5% of the stationary average $\lambda t_d/\ln(2)$. For the results shown in Fig. 5 of the main text, simulated cells that lost the prion prior to the waiting time were discarded, and the simulation were restarted until the desired number of prion containing cells to be analyzed was achieved.

#### *3.2.3. Estimating model parameters.*

**3.2.3.1 Growth rate.** Model parameters were chosen to model the experiment whose data is shown in Fig. 1f and 2 of the main text. The average doubling time for this experiment was measured to be 49.4 min. We thus set $t_d$ in our cellular growth model to 50 min. Without loss of generality we set $V_0 = 1$.

**3.2.3.2 Soluble fold production parameter.** When there are no prions, the system dynamics are governed by Eq. (1) and Eq. (4), which with $t_d = 50$ min and $V_0 = 1$, depends only on one parameter: $\lambda$. The average protein concentration for this system will always reach the stationary value of $\lambda t_d/\ln(2)$. Therefor, to determine $\lambda$ we simulated the system without prions until it reached the stationary average and then computed the average absolute partitioning error over 1000 simulations, see Fig. S14. We then set $\lambda$ such that the computed absolute partitioning error matched with that measured in our experiments when cells no longer had any prions.

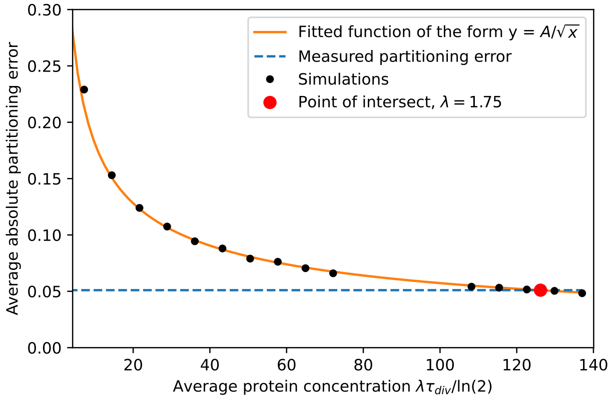

**Fig. S14. Estimating the soluble fold production parameter using partitioning errors of cells without prions.** Setting $t_d = 50$ min and $V_0 = 1$, our model admits a unique absolute partitioning error for a given $\lambda$ when there are no prions. Each black dot corresponds to the average absolute partitioning error taken over 1000 simulations. The initial condition for each simulation was set to $\lambda t_d/\ln(2)$, and each simulation ran for 1000 min of simulation time and the absolute partitioning error was recorded for the first occurring cell division after the waiting time. We fit a function of the form $y = A/\sqrt{x}$ to the simulations, with $A = 0.573$. In our experiments we measured an average absolute partitioning error for cells that have loss the prion to be 0.051. Solving for $0.051 = A/\sqrt{\lambda t_d/\ln(2)}$ gives $\lambda = 1.75$.

**3.2.3.3 Elongation and fragmentation parameters.** With $t_d = 50$ min, $V_0 = 1$, and $\lambda = 1.75$ min$^{-1}$, the remaining parameters are $\gamma$ and $\alpha$. To estimate these, we performed a parameter sweep, whereby the average time of loss, the absolute partitioning error, the average number of aggregates, and the average aggregate size was computed for different values of $\gamma$ and $\alpha$, see Fig. S15. This reveals the existence of equipotential curves in the average time of loss and absolute partitioning error plots that intersect. We thus estimated $\gamma$ and $\alpha$ by selecting the parameters of the simulated dot that minimized the following distance function: $d = \left(\frac{t_{loss}-t_{loss,m}}{t_{loss,m}}\right)^2 + \left(\frac{|P_{err}|-|P_{err,m}|}{|P_{err,m}|}\right)^2$, where $t_{loss}$ and $|P_{err}|$ are the average time of loss and absolute partitioning error respectively, and $t_{loss,m} = 129.22$ min and $|P_{err,m}| = 0.125$ are the experimental measurements. This gives us an estimate of $\gamma = 1.122 \times 10^{-3}$ min$^{-1}$ and $\alpha = 1.970 \times 10^{-4}$ min$^{-1}$, indicated by the orange dot in Fig. S15.

To make the contour plots shown in Fig. 5 of the main text, we binned the log-log plots shown in Fig. S15 into a 16x16 grid with bins that scaled logarithmically, and computed the averages in each bin to eliminate sampling noise. The python function tricontour taken from matplotlib was used on the binned averages to generate the contour plots.

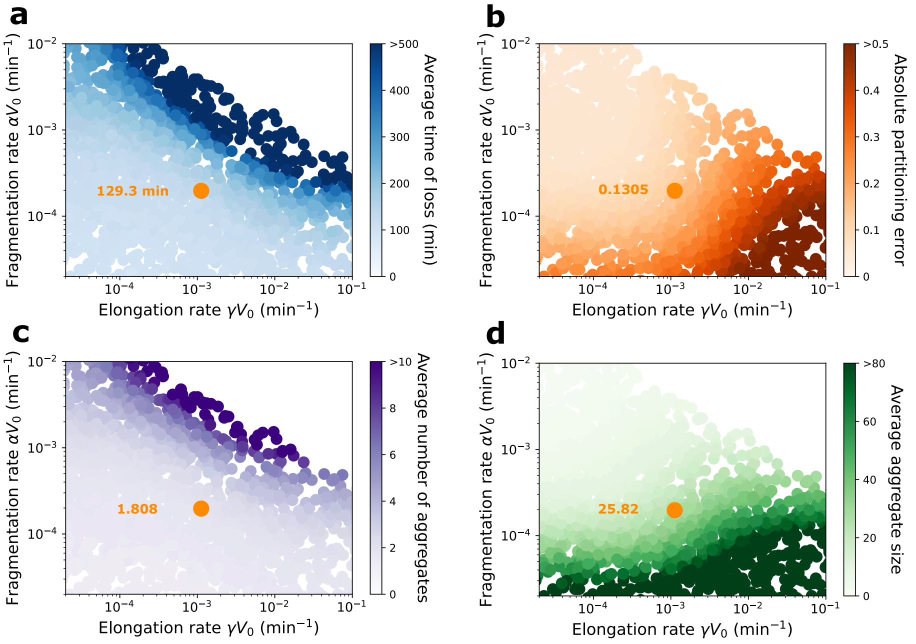

**Fig. S15. Estimating the elongation and fragmentation parameters using partitioning errors of cells with prions and their time of prion loss.** Each dot corresponds to an average taken over 1000 simulations, each with the initial conditions described in Sec. 3.2.2.2., with $t_d = 50$ min, $V_0 = 1$, and $\lambda = 1.75$ min$^{-1}$. Analysis started after the waiting time described in Sec. 3.2.2.3. The orange dot and text corresponds to the set of parameters simulated that most closely matched with the experimentally measured time of loss and absolute partitioning error values. Simulations were not performed in the top right region because simulations in this region are computationally time consuming. **a)** The average time of loss grows as the elongation and fragmentation rates grow. Time of loss is defined as the time when $Y_k = 0$ for all $k$. If the prions were all lost prior to the waiting time, the simulation was discarded and a new one was started until 1000 times of loss were computed. **b)** The absolute partitioning error prior to loss grows as the elongation rate grows. The absolute partitioning error was taken as the average over all divisions that occurred in the 1000 simulations prior to and at prion loss. **c)** The average number of aggregates grows similarly to the average time of loss. This is because uneven partitioning of molecules at cell division decreases as the number of molecules increases. Here the average number of aggregates was taken over the 1000 time trajectories prior to prion loss, computed as $\langle Y_{tot} \rangle = \sum_i i \times$ (Total time spent when $Y_{tot} = i$) / (Total time spent over all simulations), where $Y_{tot} = \sum_k y_k$. **d)** The average aggregate size grows similarly to the absolute partitioning error. Here the average aggregate size was taken over all trajectories prior to prion loss, computed as $\sum_k k\langle Y_k \rangle / \langle Y_{tot} \rangle$, with $\langle Y_k \rangle = \sum_i i \times$ (Total time spent when $y_k = i$) / (Total time spent over all simulations).

***3.2.4. Different initial conditions.*** We used the stochastic model to see whether the initial prion concentration could affect the time-of-loss curves. In particular, with the parameters estimated in the previous sections, we computed the time-of-loss curves for three different systems that differ by their initial prion concentration, see Fig. S16. All three systems start with one large aggregate of different size. We find that the time-of-loss curves are identical.

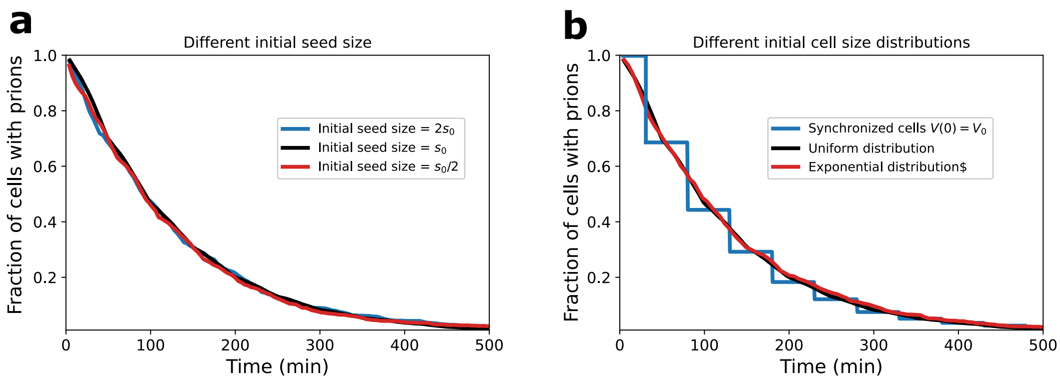

**Fig. S16. Prion loss kinetics are unaffected by different initial seed sizes in the stochastic model a)** Time of loss curves were computed for three systems with different initial seed sizes. In Sec. 3.2.2.2 we describe how we estimated an initial seed size of $s_0 = \lambda t_d V(0)/\ln(2)$. We then computed time-of-loss curves where the initial seed size is $2s_0$ and $s_0/2$. The time-of-loss curves are identical. Each curve corresponds to 1000 simulations. **b)** Time of loss curves were computed for three systems with different initial cell size distributions. First we set the cell size distribution to be a uniform distribution that ranges from $V_0$ to $2V_0$. Next we set the distribution to be an exponential with a probability density function of the form $\text{prob}(V) = 2\ln(2)^2 e^{-\ln(2)V}$. Lastly we looked at the case where each cell is synchronized and starts with the same initial cell size $V(t=0) = V_0$. The time-of-loss curves for the uniform and exponential distributions are identical. The time-of-loss curve for the synchronized cells follows steps with intervals given by $t_d$ because the division times are all synchronized.

Moreover, so far in our simulations each cell starts with an initial cell size that is picked from a uniform distribution that ranges from $V_0$ to $2V_0$. In the mother machine experiments cells are first grown in a growing cell population to stationary phase which could lead to a non-uniform distribution of initial cell sizes. To see whether a non-uniform initial cell size distribution would affect the observed time-of-loss curves, we computed these curves for three systems with different initial cell size distributions, see Fig. S16. We find that the three curves are identical.

***3.2.5. Non-zero minimum seed size.*** In the previous sections, the minimum seed size was set to zero. Next, we set the initial seed size to 2, where any prion aggregate of size 2 or less immediately breaks apart into soluble fold proteins. We find that in spite of this change the estimated elongation and fragmentation rates are similar to the zero minimum seed size case, see Fig. S17.

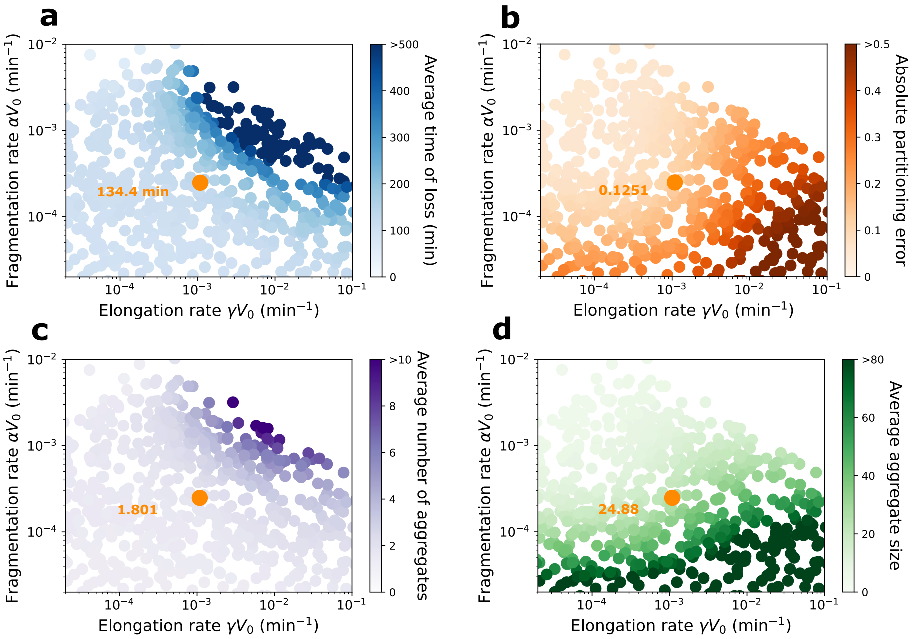

**Fig. S17. A minimum seed size of two has little effect on the estimated elongation and fragmentation parameters.** Each dot corresponds to an average taken over 1000 simulations, each with the initial conditions described in Sec. 3.2.2.2, with $t_d = 50$ min, $V_0 = 1$, and $\lambda = 1.75$ min$^{-1}$. Analysis started after the waiting time described in Sec. 3.2.2.3. The orange dot and text corresponds to the set of parameters simulated that most closely matched with the experimentally measured time of loss and absolute partitioning error values. Simulations were not performed in the top right region because simulations in this region are computationally time consuming. **a)** The average time of loss grows similarly to that in Fig. S15a, except in the top left corner where it does not grow. This is because a high fragmentation rate means smaller aggregates. Eventually small aggregates become close to the minimum seed size where they are converted out of prion form which can drive prion loss along with cell division. **b)** The absolute partitioning error prior to loss grows as the elongation rate grows. **c)** The average number of aggregates grows similarly to the average time of loss. Here the average number of aggregates was taken over the 1000 time trajectories prior to prion loss, computed as $\langle Y_{tot} \rangle = \sum_i i \times$ (Total time spent when $Y_{tot} = i$) / (Total time spent over all simulations), where $Y_{tot} = \sum_k y_k$. **d)** The average aggregate size grows similarly to the absolute partitioning error. Here the average aggregate size was taken over all trajectories prior to prion loss, computed as $\sum_k k\langle Y_k \rangle / \langle Y_{tot} \rangle$, with $\langle Y_k \rangle = \sum_i i \times$ (Total time spent when $y_k = i$) / (Total time spent over all simulations).

***3.2.6. Estimating the replication rate.*** Here we derive an expression for the replication rate (defined in (13)) in terms of the parameters of the stochastic model shown in Fig. 5a of the main text and defined in Sec. 3.2.1.

In Supplementary Note 1 of (13), the following equations model the growth of aggregates in the absence of aggregate removal

$$\frac{dM}{dt} = k_g G(t) \quad \& \quad \frac{dG}{dt} = k_m M(t), \qquad [9]$$

where $M(t)$ is the total aggregated mass, $G(t)$ is the concentration of growth-competent ends in linear aggregation, $k_g$ is the rate of growth, and $k_m$ is the rate of multiplication. The replication rate is then defined as $\kappa = \sqrt{k_g k_m}$.

In terms of the stochastic model defined in Sec. 3.2.1, the total aggregated mass is given by $M(t) = m\sum_{k=1}^{\infty} k y_k$, where $m$

is the mass of one monomer. In the regime where there is no removal of aggregates (i.e., when there is no cell division), the system is driven solely by the reactions given by Eqs. (1), (2), and (3). Note that $M(t)$ remains unchanged when the reactions of Eq. (1) and Eq. (3) occur because the total number of monomers that make up aggregates remains unchanged. Therefore, $M(t)$ increases only when elongation occurs:

$$(x,\ M(t)) \xrightarrow{r_{\text{elon}}} (x-1,\ M(t)+m)\,,$$

where the total elongation rate is $r_{\text{elon}} = \gamma\,[X]\,[Y_{\text{tot}}]\,V$ with $[Y_{\text{tot}}] = \sum_k [Y_k] = \sum_k y_k/V$. In the deterministic limit this becomes

$$\frac{dM}{dt} = m\gamma\,[X]\,[Y_{\text{tot}}]\,V. \qquad [10]$$

Since $G(t)$ is defined as the concentration of growth competent sites, we have $G(t) = 2\,[Y_{\text{tot}}]$ since there are two sites that a monomer can bind to a linear aggregate. Eq. (10) then becomes

$$\frac{dM}{dt} = k_g G(t), \qquad [11]$$

where $k_g = m\gamma\,[X]\,V/2$ is the rate of growth.

Next, we note that the concentration of total aggregates does not change when reactions given by Eq. (1) and Eq. (2) occur. Therefore, $G(t)$ increases only when fragmentation occurs:

$$G(t) \xrightarrow{r_{\text{frag}}} G(t) + 2/V,$$

where the total fragmentation rate is $r_{\text{frag}} = \sum_{k=1}^{\infty}(k-1)\alpha\,[Y_k]\,V$ because each aggregate of size $k$ has $(k-1)$ bindings that can break. Note that the reaction step size is $2/V$ because when an aggregate fragments there is an increase of two aggregate sites from each end of the new aggregate, and so the total *concentration* increases by $2/V$. In the deterministic limit this becomes

$$\frac{dG}{dt} = \frac{2}{V}\sum_k (k-1)\alpha\,[Y_k]\,V = \frac{2}{V}\left(\sum_{k=1}^{\infty} k\alpha\,[Y_k]\,V - \sum_{k=1}^{\infty}\alpha\,[Y_k]\,V\right). \qquad [12]$$

Since the average aggregate size in our parameter regime is $\approx 25$ (see Fig. S15d), we make the approximation that the second term on the right is negligible compared to the first.

$$\Rightarrow \frac{dG}{dt} \approx \frac{2}{V}\sum_{k=1}^{\infty} k\alpha\,[Y_k]\,V = \frac{2\alpha}{mV}\cdot m\sum_{k=1}^{\infty} k y_k = k_m M(t), \qquad [13]$$

where $k_m = 2\alpha/mV$ is the rate of multiplication. Comparing Eqs. (11) and (13) with Eq. (10), the replication rate is given by $\kappa = \sqrt{k_g k_m} = \sqrt{[X]\,\alpha\gamma}$. Substituting the estimated $\alpha$ and $\gamma$ from Sec. 3.2.3.3, along with an average monomer concentration of approximately 100 as shown in Fig. S10d, gives us $\kappa \approx 7.84 \times 10^{-5} s^{-1} \sim 10^{-5} s^{-1}$.
